# Distinct regions of the kinesin-5 C-terminal tail are essential for mitotic spindle midzone localization and sliding force

**DOI:** 10.1101/2023.05.01.538972

**Authors:** Zachary R. Gergely, Michele H. Jones, Bojun Zhou, Cai Cash, J. Richard McIntosh, Meredith D. Betterton

**Affiliations:** Department of Physics, University of Colorado Boulder, Boulder CO 80309; Department of Molecular, Cellular, and Developmental Biology, University of Colorado Boulder, Boulder CO 80309

## Abstract

Kinesin-5 motor proteins play essential roles during mitosis in most organisms. Their tetrameric structure and plus-end-directed motility allow them to bind to and move along antiparallel microtubules, thereby pushing spindle poles apart to assemble a bipolar spindle. Recent work has shown that the C-terminal tail is particularly important to kinesin-5 function: the tail affects motor domain structure, ATP hydrolysis, motility, clustering, and sliding force measured for purified motors, as well as motility, clustering, and spindle assembly in cells. Because previous work has focused on presence or absence of the entire tail, the functionally important regions of the tail remain to be identified. We have therefore characterized a series of kinesin-5/Cut7 tail truncation alleles in fission yeast. Partial truncation causes mitotic defects and temperature-sensitive growth, while further truncation that removes the conserved BimC motif is lethal. We compared the sliding force generated by *cut7* mutants using a kinesin-14 mutant background in which some microtubules detach from the spindle poles and are pushed into the nuclear envelope. These Cut7-driven protrusions decreased as more of the tail was truncated, and the most severe truncations produced no observable protrusions. Our observations suggest that the C-terminal tail of Cut7p contributes to both sliding force and midzone localization. In the context of sequential tail truncation, the BimC motif and adjacent C-terminal amino acids are particularly important for sliding force. In addition, moderate tail truncation increases midzone localization, but further truncation of residues N-terminal to the BimC motif decreases midzone localization.

---

Kinesin-5 motors are essential for mitosis in many organisms because they help assemble the bipolar mitotic spindle (Fig. 1) [1–5]. Kinesin-5 tetramers are formed by two dimers linked in antiparallel orientation through their stalks, so the motor heads are positioned at both ends of a bipolar complex [6–11]. When kinesin-5s crosslink antiparallel microtubules (MTs), their plus-end-directed motility generates force that slides the microtubules apart [12–18]. This sliding force is important for initial separation of mitotic spindle poles, assembly of a bipolar spindle (Fig. 1A), and for spindle elongation in early anaphase B [19–21]. Loss of kinesin-5 function commonly leads to the formation of monopolar mitotic spindles [2, 4, 22], consistent with the role of kinesin-5 sliding force in separating spindle poles.

**FIG. 1.**
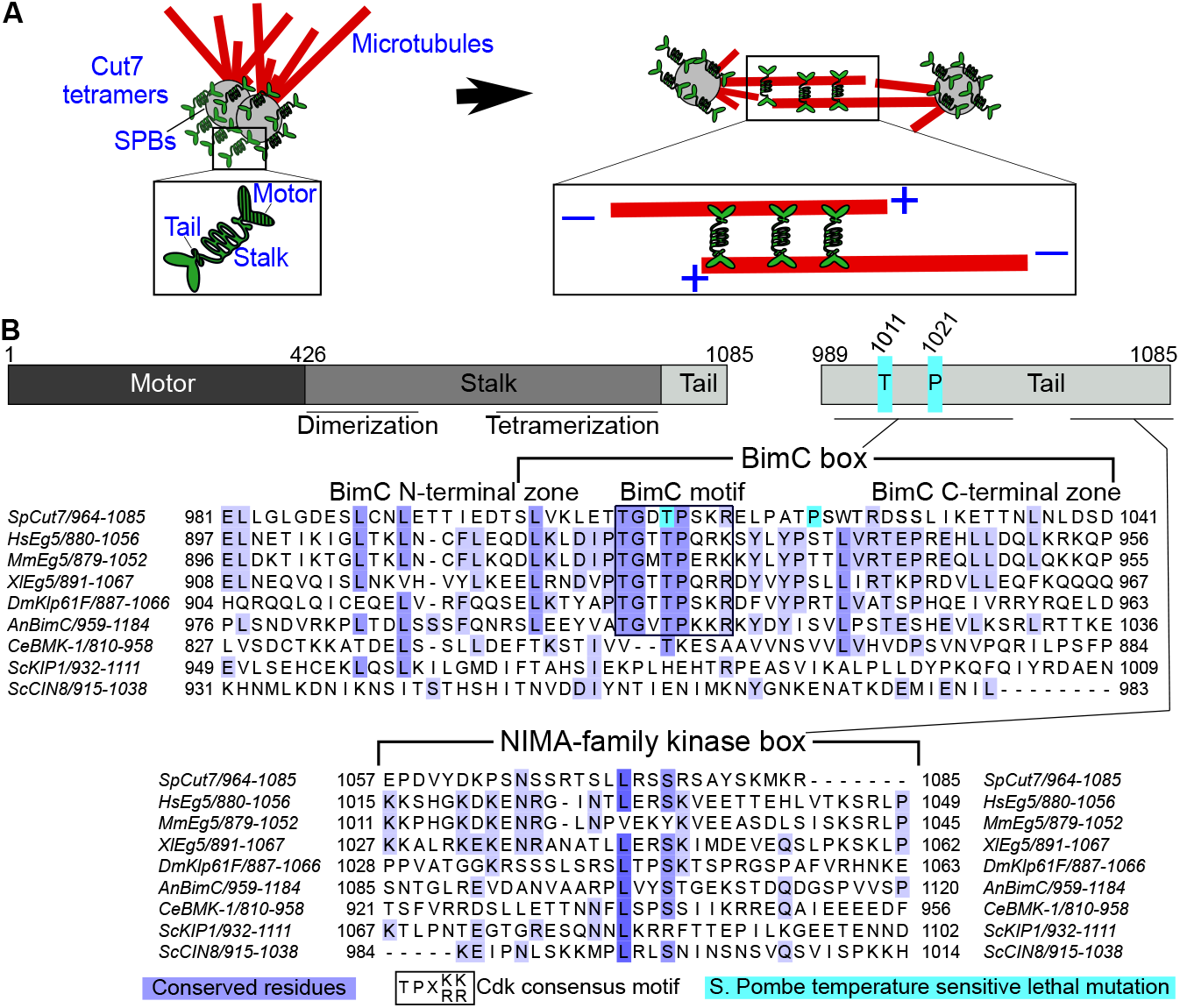
Overview of kinesin-5 mitotic function and C-terminal tail. (A) Schematic of Cut7p and its role in bipolar spindle assembly. Left: Cut7p tetramers near the fission-yeast spindle-pole bodies (SPBs) of a monopolar spindle. Inset: Cut7p tetramer showing MT-binding motor, stalk with dimerization and tetramerization regions, and C-terminal tail. Right: Cut7p tetramers on a bipolar spindle. Inset: Cut7p crosslinking antiparallel microtubules in the spindle midzone. (B) Upper: domain map of *cut7* showing the motor, stalk, and ∼ 96-amino acid C-terminal tail. Middle and lower: ClustalW sequence alignment of kinesin-5 C-terminal tails near the BimC box (middle) and NIMA-family kinase box (lower), adapted from Ref. [47]. Alignment includes *Schizosaccharomyces pombe cut7, Homo sapiens* Eg5/KIF11, *Mus musculus* Eg5, *Xenopus laevis* Eg5, *Drosophila melanogaster* KLP61F, *Aspergillus nidulans* BimC, *Caenorhabditis elegans* BMK-1, and *Saccharomyces cerevisiae* Kip1 and Cin8. Conserved residues are highlighted in purple, *S. pombe* temperature-sensitive lethal mutations are highlighted in cyan, and the Cdk consensus motif is boxed [5, 42, 45–47, 55, 56].

However, increasing evidence suggests that this model for how kinesin-5s separate spindle poles is incomplete. Kinesin-5s are not only plus-end-directed, but can also walk or be carried toward spindle MT minus ends. Fungal kinesin-5s have intrinsic bidirectional motility [23–34], while in vertebrates, kinesin-5 becomes cargo for a minus-end directed motor [35–39]. At least in part because of this minus-end-directed motility, kinesin-5s localize more strongly near spindle poles (near MT minus ends) than in the center of the spindle, where MT plus ends predominate [4, 5, 38, 40–43]. While the functional significance of kinesin-5 minus-end-directed motility and pole localization have remained unclear, previously work suggested that pole localization may position kinesin-5s for spindle elongation [32, 44]. More recently, we proposed that these mechanisms may sequester kinesin-5 near the poles to limit sliding force [34]. In this light, the mechanisms by which kinesin-5 directionality and localization are controlled may be important to its regulation.

Kinesin-5 monomers have a conserved organization with an N-terminal motor domain, a stalk containing oligomerization domains, and at the C-terminus of the protein, ∼96 amino acids predicted to be disordered, known as the C-terminal tail (Fig. 1B). While the greatest amino acid conservation is in the motor domain, the tail has conserved regions known as the BimC box and NIMA-family kinase box [5, 41, 45–47]. These include consensus sites for cyclin-dependent kinase 1 (Cdk) and NIMA-family kinase (NFK, Fig. 1B). Because kinesin-5s are antiparallel tetramers, the C-terminal tail domain of one dimer lies near the motor domain of the other dimer (Fig. 1A). This geometry has led to a recent focus on the possible role of tail-motor interactions in kinesin-5 motility. It has been observed that cancer cells can develop resistance to kinesin-5/Eg5 inhibitors via tail truncation[48], further supporting the significance of the tail.

The kinesin-5 tail regulates motor function and localization. For metazoan kinesin-5s, truncation of the C-terminal tail alters motility, MT crosslinking, and sliding force of purified motor [18, 49, 50]. However, because the tail domain is also critical for spindle localization [18], the importance of the tail for kinesin-5 force in the spindle cannot be assessed for these organisms. Closed mitosis in fungi appears to have advantages in confining kinesin-5 tail truncations in a smaller volume where they can maintain spindle localization. Deletion of the kinesin-5/Cin8 tail in budding yeast biases directionality toward MT plus ends [51] and is lethal if the gene for a second kinesin-5 allele (Kip1) is also deleted [14, 51]. In fission yeast, truncation of the kinesin-5/Cut7 tail is lethal and affects bipolar spindle assembly [34]. Notably, however, *cut7* becomes inessential in fission-yeast lacking kinesin-14 [52–54]. We used this genetic background to show that tail truncation alters Cut7p velocity and rate of direction switching [34]. While the C-terminal tail is predicted to be disordered based on its primary structure [34], cryo-EM analysis of *Drosophila* KLP61F found that interactions between the tail and motor domains can at least partially fold the tail [18]. Tail-motor interactions also alter motor structure, causing slower ATP hydrolysis and speed of purified motor [18]. Further, it has recently been proposed that C-terminal tails of kinesin-5 can interact with motor domains of adjacent motors, promoting cluster formation that enhances sliding force [50] and could affect direction switching [30, 33]. However, the extent and mechanisms of kinesin-5 regulation by the C-terminal tail remains incompletely understood.

The fission yeast *Schizosaccharomyces pombe* is a useful model in which to study kinesin-5. Cut7p is the sole kinesin-5 motor in this organism, and it shares domain structure and spindle localization with other kinesin-5s (Fig. 1). The Cut7p motor-domain structure has been solved [57]. In fission yeast lacking kinesin-14/Pkl1 or its binding partners, Wdr8p and Msd1p, spindle MT minus ends can be pushed past the spindle-pole body (SPB) and distort the nuclear envelope outward [52–54]. These spindle MT protrusions can differ in length, dynamics and in the polarity of the MTs they contain [54]. Importantly, spindle MT minus-end protrusions depend on Cut7p, since they do not form in cells in which Cut7 is deleted or has a loss of function [53, 54]. Protrusions can therefore assay the net Cut7p force, determined by the balance of spindle elongation force with drag from Cut7p crosslinking.

## RESULTS

### Sequential truncation of the *cut7* C-terminal tail leads to growth and spindle defects

In previous work, we found that the *cut7* tail truncation lacking the C-terminal 96 amino acids (*cut7-988*) was lethal in an otherwise wild-type genetic background [34]. To study which portions of the *cut7* tail are most important for cell viability, spindle assembly, and net sliding force, we generated a series of sequential C-terminal truncation mutants as well as an internal deletion (Fig. 2A). We constructed each *cut7* allele with a green fluorescent protein (GFP) tag at its C-terminus and integrated it as the sole copy of *cut7* in the cell. In addition, we added low-level-expressed *mCherry-atb2* to label spindle MTs [34, 58, 59].

**FIG. 2.**
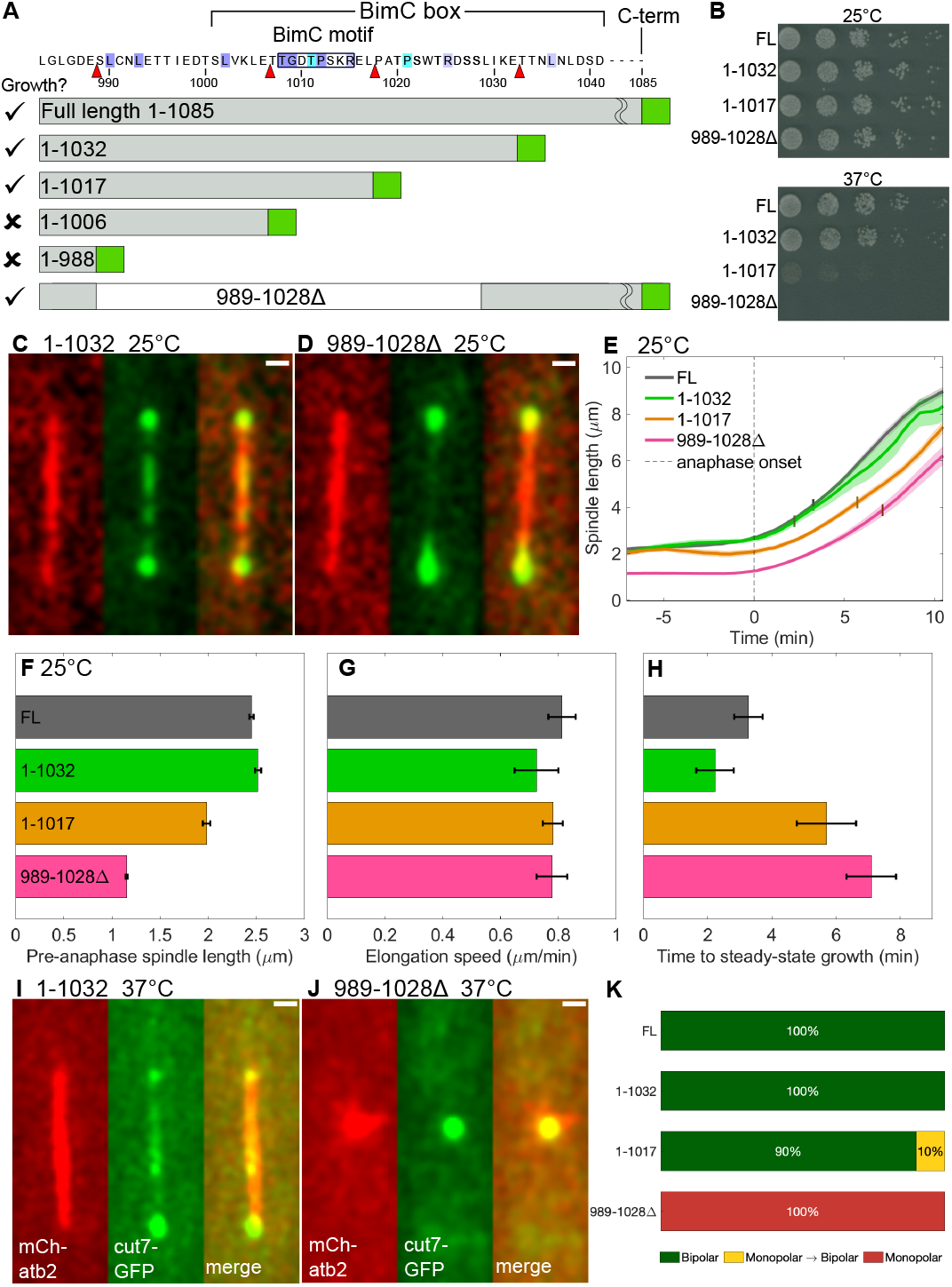
Sequential truncation of the *cut7* C-terminal tail compromised cell viability and spindle assembly and function. (A) Schematic of truncation mutants aligned with the BimC box region amino acid sequence (upper), showing conserved residues (purple), temperature-sensitive lethal mutations (cyan) and truncation sites (red triangles). Each allele is labeled with amino acids retained (gray bar and numbers), C-terminal GFP (green bar), and viability (left column). (B) Serial dilution assays of cells grown at 25°C (upper) and 37°C (lower). All viable strains grew at 25°C, and *cut7-1017* and *cut7-989-1028*Δ showed temperature-sensitive growth inhibition or lethality. (C-D) Fluorescence images showing mCherry-atb2 MTs (red, left), Cut7-GFP (green, center), and merge (right) for cells grown and imaged at 25°C and shown as maximum-intensity projections of confocal sections. (C) Cut7-1032-GFP localized near SPBs and was visible on the interpolar spindle. (D) Cut7-989-1028Δ-GFP localized near SPBs and appeared dimmer at the spindle midzone. Scalebar, 1 *µ*m. (E) Spindle length dynamics. Spindle length as a function of time was averaged for 19 *cut7*, 9 *cut7-1032*, 11 *cut7-1017*, and 14 *cut7-989-1028*Δ cells and aligned with *t* = 0 the time of anaphase onset (dashed vertical line, Methods). Time at which steady anaphase elongation speed was reached marked on each curve by the vertical bar. (F) Pre-anaphase spindle length, (G) anaphase spindle elongation speed, and (H) the time between anaphase onset and start of steady elongation. Error bars in E-H are standard error of the mean. (I-J) Fluorescence images showing mCherry-atb2 MTs (red, left), Cut7-GFP (green, center), and merge (right) for cells grown and imaged at 37°C and shown as maximum-intensity projections of confocal sections. (I) Cut7-1032-GFP localizes similarly to Cut7-GFP on the bipolar spindle. (J) Cut7-989-1028Δ-GFP localizes near the center of the monopolar spindle due to a failure of bipolar spindle assembly. Scalebar, 1 *µ*m. (K) Fraction of cells grown at 37°C that formed bipolar spindles (green), monopolar spindles that became bipolar (yellow), or persistent monopolar spindles (red) for 21 *cut7*, 17 *cut7-1032*, 21 *cut7-1017*, and 21 *cut7-989-1028*Δ cells.

While truncation of small portions of the tail did not noticeably affect cell growth, greater truncation caused temperature-sensitive growth defects and lethality. The *cut7-1032* truncation removed the final 53 amino acids from the tail, including the NIMA-family kinase box. Growth was similar for *cut7-1032* and full-length *cut7* cells (*cut7-FL*, Fig. 2B). Cells with *cut7-1017* lacking the final 68 amino acids grew similarly to *cut7-FL* and *cut7-1032* at 25°C, but showed a strong growth defect at 37°C. The *cut7-1006* allele lacked 79 amino acids, including the conserved BimC motif (Fig. 2A). We were unable to obtain *cut7-1006* cells by transformation as the sole *cut7* allele in otherwise wild-type cells, suggesting it was lethal. The *cut7-988* mutant lacked the entire tail up to the predicted coiled-coil sequence, and was a lethal allele [34]. Finally we made an internal deletion of a.a. 989-1028, which lacked the BimC motif, but retained a C-terminal portion of the BimC box and the NIMA-family kinase box (Fig. 2A) [47]. Cells with *cut7-989-1028*Δ showed a slight growth defect at 25°C and were inviable at 37°C (Fig. 2B, S6). These results suggest that the BimC box and nearby residues are particularly important for cell growth.

The viable truncation mutants all showed bipolar spindle assembly at 25°C with spindle-localized Cut7-GFP, but differed in spindle length and elongation (Figs. 2C-E, S1, S6). We imaged cells by confocal fluorescence microscopy, then quantified spindle length over time and aligned traces by defining *t* = 0 as the time of anaphase onset (Methods, Fig. S1). We measured pre-anaphase spindle length, anaphase spindle elongation speed, and the time from anaphase onset until steady spindle elongation was reached (Fig. 2E-H, Methods). The *cut7-FL* and *cut7-1032* spindles appeared similar, with pre-anaphase spindle length of ∼2.5 *µ*m, anaphase spindle elongation speed of ∼ 0.8 *µ*m/min, and a ∼2-3 min time after anaphase onset to reach steady elongation (Fig. 2E-H). Cells containing *cut7-1017* and *cut7-989-1028*Δ had shorter pre-anaphase spindles (2 and 1.25 *µ*m, Fig. 2F). While the steady anaphase elongation speed was similar in all strains (Fig. 2G), *cut7-1017* and *cut7-989-1028*Δ cells took longer to reach steady elongation (Fig. 2H). This suggests that Cut7p with the tail truncated at a.a. 1017 or with the internal deletion might have defects in Cut7p net sliding force needed to elongate the spindle.

Temperature-sensitive growth defects correlated with spindle defects in truncation mutants. All Cut7-GFP mutants localized to the spindle at 37°C in confocal fluorescence microscopy (Fig. 2I,J). All *cut7-FL* and *cut7-1032* cells imaged assembled a bipolar spindle (Fig. 2K). By contrast, *cut7-1017* cells, which showed growth defects at 37°C, also showed spindle assembly delays in 10% of cells (Fig. 2K), suggesting that spindle defects caused the growth defects. For *cut7-989-1028*Δ cells that were inviable at restrictive temperature, all cells observed showed spindle assembly failure with monopolar spindles (Fig. 2J,K). Of the viable mutants we studied, *cut7-989-1028*Δ cells with the BimC motif removed had the most severe phenotype (Fig. S6).

### The BimC motif is required for proper Cut7p spindle midzone localization

Previous work showed that *cut7-988* lacking the entire tail showed changes in Cut7p localization and motility, suggesting that a reduced number of motors in the spindle midzone led to lower net sliding force [34]. Therefore, we asked whether localization or motility changes might explain *cut7* truncation mutant phenotypes at permissive temperature. We used automated image analysis (Methods, [34, 60]) to create kymographs of Cut7-GFP intensity along the spindle axis, determine average intensity profiles, and compute both total Cut7-GFP spindle fluorescence and the ratio of intensity at the midzone to that at the poles (Fig. 3, Methods). Because Cut7-GFP shows enhanced localization at spindle poles [34, 40, 52, 61] (Fig. 3A), we grouped profiles of Cut7-GFP fluorescence by peak-to-peak distance and averaged (Fig. 3B-D). This allowed us to measure how Cut7-GFP fluorescence redistributes as spindle length changes during mitosis. Cut7-FL-GFP average fluorescence intensity suggested that as the spindle increased in length from 2 to 4 *µ*m, the midzone intensity increased slightly (Fig. 3B,C, black lines). Once spindle length reached 7 *µ*m, the midzone intensity decreased and pole intensity became asymmetric (Fig. 3D, Methods). This quantitative analysis was consistent with previous [34, 40, 52, 61] and current (Fig. 3A) qualitative results.

**FIG. 3.**
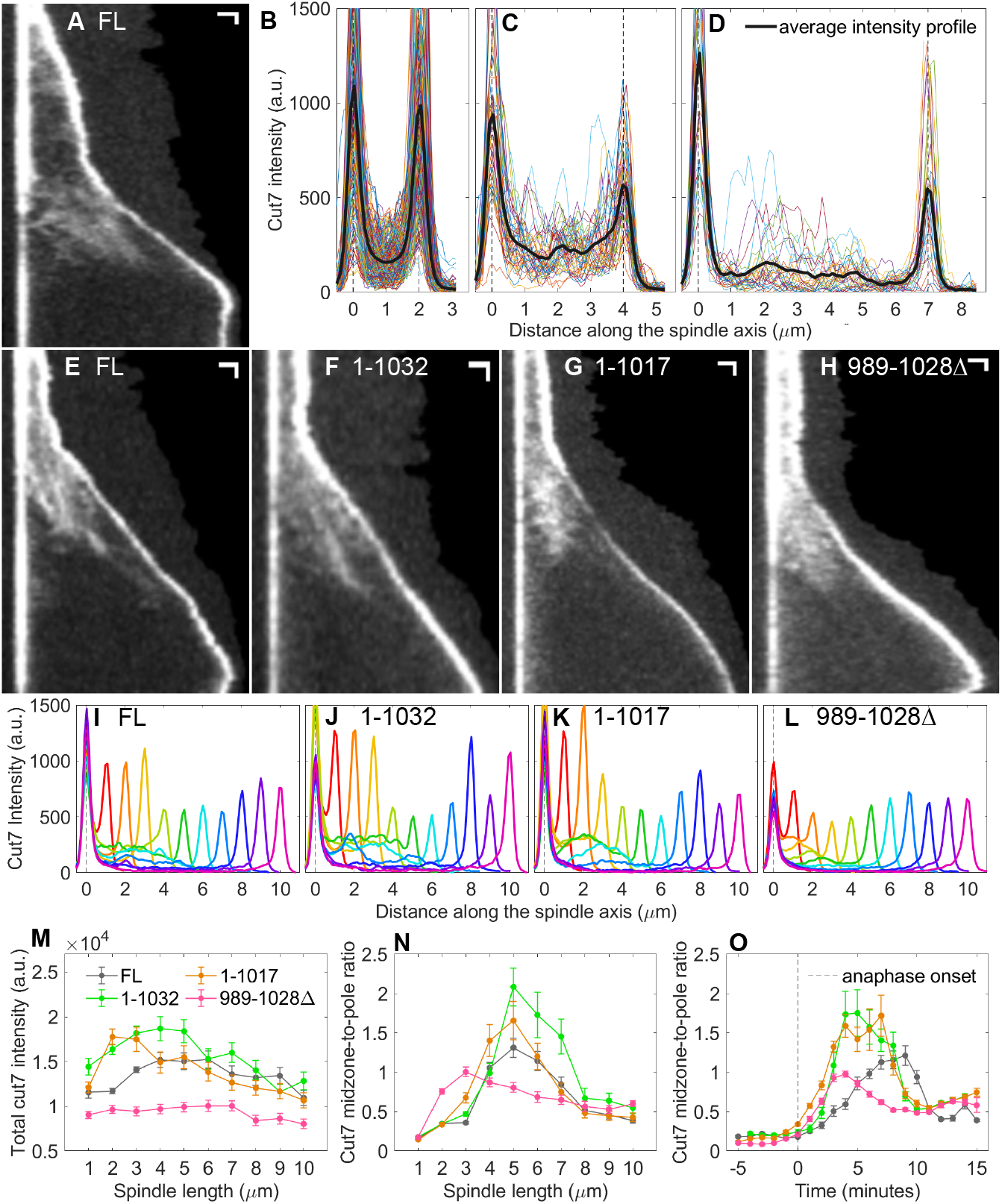
Tail truncated Cut7-GFP mutants bound to and moved along spindle MTs. (A) Kymograph of Cut7-GFP movement on a bipolar spindle. Bright regions show SPB-localized signal and dim streaks show midzone-localized moving clusters. Scalebars, 1 *µ*m horizontal, 60 sec vertical. (B-D) Cut7-GFP intensity along the spindle axis for spindle length of 2 *µ*m (B), 4 *µ*m (C) and 7 *µ*m (D). Colored lines show data from individual images, black line shows average. Data from 202 images (B), 45 images (C), and 28 images (D), from 29 cells. (E-H) Kymographs of Cut7-FL-GFP (E), Cut7-1032-GFP (F), Cut7-1017-GFP (G), and Cut7-989-1028Δ-GFP (H). (I-L) Average Cut7-GFP intensity along the spindle axis for Cut7-FL-GFP (I), Cut7-1032-GFP (J), Cut7-1017-GFP (K), and Cut7-989-1028Δ-GFP (L). (M) Total Cut7-GFP intensity on the spindle as a function of spindle length. (N) Cut7-GFP midzone-to-pole ratio as a function of spindle length. (O) Cut7-GFP midzone-to-pole ratio as a function of time relative to anaphase onset (*t* = 0, dashed vertical line). Intensity data from 29 *cut7*, 34 *cut7-1032*, 29 *cut7-1017*, and 30 *cut7-989-1028*Δ cells. Error bars in M and N are the standard error of the mean assuming independent intensity measurements for each image at each integer spindle length. Error bars in O are the standard error of the mean assuming independent intensity measurements are separated by 1 minute time intervals.

Kymographs of Cut7-GFP movement on the spindle appeared similar for all the viable truncation mutants (Fig. **??**A,E-H]fig:trunc-data). Therefore, at permissive temperature, truncation mutants of *cut7* localize to and move on spindle MTs. This suggests that the spindle length and elongation defects we observed do not occur due to complete loss of Cut7p motility or tetramerization. However, upon inspection, Cut7-989-1028Δ-GFP kymographs appeared to have lower overall intensity and lower relative midzone intensity. Therefore, we quantified subtle changes in localization for different Cut7-GFP alleles, by averaging intensity profiles over a range of spindle length (Fig. 3I-L, S2, S3, S4, S5, Methods). While the average intensity profiles appeared similar for the full-length motor and truncation mutants, Cut7-989-1028Δ-GFP showed lower intensity, both at the spindle midzone and at poles. To quantify fluorescence, we computed the total Cut7-GFP intensity on the spindle for varying spindle length (Fig. 3M). Cut7-989-1028Δ-GFP indeed showed lower total intensity than the other strains. Lower Cut7p expression, decreased stability, or a spindle localization defect in these cells could lead to lower amounts of motor on the spindle. Therefore, even if Cut7-989-1028Δ-GFP produced the same sliding force per motor, decreased motor levels could lead to shorter spindles and longer time before steady anaphase elongation in this mutant (Fig. 2E-H).

Removal of the BimC motif also altered the balance of Cut7p at the spindle midzone compared to the poles. Recently, we proposed that spindle poles may serve as a “waiting room” where kinesin-5s are sequestered, lowering motor number to limit sliding force in the midzone [34]. Since our observations of altered spindle length and steady anaphase elongation delays suggested alterations in net Cut7p sliding force in some mutants, we asked whether this might be caused by shifts in Cut7p distribution along the spindle. We therefore quantified the ratio of midzone-to-pole Cut7-GFP fluorescence (Methods). Because fission yeast contain overlapping antiparallel MTs throughout the interpolar spindle [62–64], we defined the spindle midzone as all spindle MTs between the poles. The average midzone-to-pole intensity ratio was low (*<*1) for short spindles, then increased (to ∼1.5) in early anaphase B, and finally decreased in late mitosis (Fig. 3N). However, Cut7-989-1028Δ-GFP showed increased midzone localization for shorter spindles than the other strains, while the overall ratio was lower for all spindle lengths observed. We additionally analyzed the midzone-to-pole ratio as a function of time relative to anaphase onset, and found that the time of increase was similar in all strains (Fig. 3O). To compare overall midzone localization, we averaged the Cut7-GFP midzone-to-pole ratio (Fig. 3N) for all spindles with length from 3 to 7 *µ*m. This gave a single average midzone-to-pole ratio that was *<* 1 only for Cut7-989-1028Δ-GFP (Fig. S6). The moderate truncation of *cut7-1032* and *cut7-1017* cells led to an increased average midzone-to-pole ratio in these cells (Fig. S6). This suggests that even if tail truncation lowered the sliding force per motor, the increased midzone protein levels could partially rescue the effects of lower force. For the internal deletion, total Cut7-989-1028Δ-GFP fluorescence was lower and the relative amount of midzone-localized motor was lower in anaphase. Both of these changes could contribute to reduced net sliding force and spindle length and elongation defects in these cells.

### Deletion of kinesin-14 motor genes rescues lethality of Cut7 tail truncation

Deletion of kinesin-14/Pkl1 suppresses the deleterious effects of *cut7* loss-of-function alleles [43, 65–68]. Since deletion of *pkl1* allowed study of the otherwise lethal *cut7-988* truncation [34], we introduced *cut7-FL* and our four truncation alleles (Fig. 2A) into the *pkl1*Δ *klp2*Δ background to compare their phenotypes. All of our strains were viable (Fig. 4A). While no growth defects occurred at 25°C for *cut7-1032* and *cut7-1017* (Fig. 4A, top), removal of the BimC motif in *cut7-1006* severely suppressed growth. We were surprised to find that removal of the entire tail in *cut7-988* showed a lesser growth defect; however, it may act more like a *cut7*Δ allele which has been shown to grow relatively better than certain *cut7* mutant alleles in this background [66, 67]. At 37°C, *cut7-1032* and *cut7-1017* grew similarly to full length, whereas *cut7-1006* was lethal and *cut7-988* had a moderate growth defect (Fig. 4A, bottom). At 37°C, the truncations that grew well showed nearly all bipolar spindles in live-cell imaging, while *cut7-1006* and *cut7-988* had monopolar spindles (Fig. S7), consistent with the growth defect occurring due to spindle assembly failure. The rescue of these previously lethal alleles allowed us to further assess the potential effects of the amino acids N-terminal to the BimC motif (Fig. 1B) on Cut7p localization and force.

**FIG. 4.**
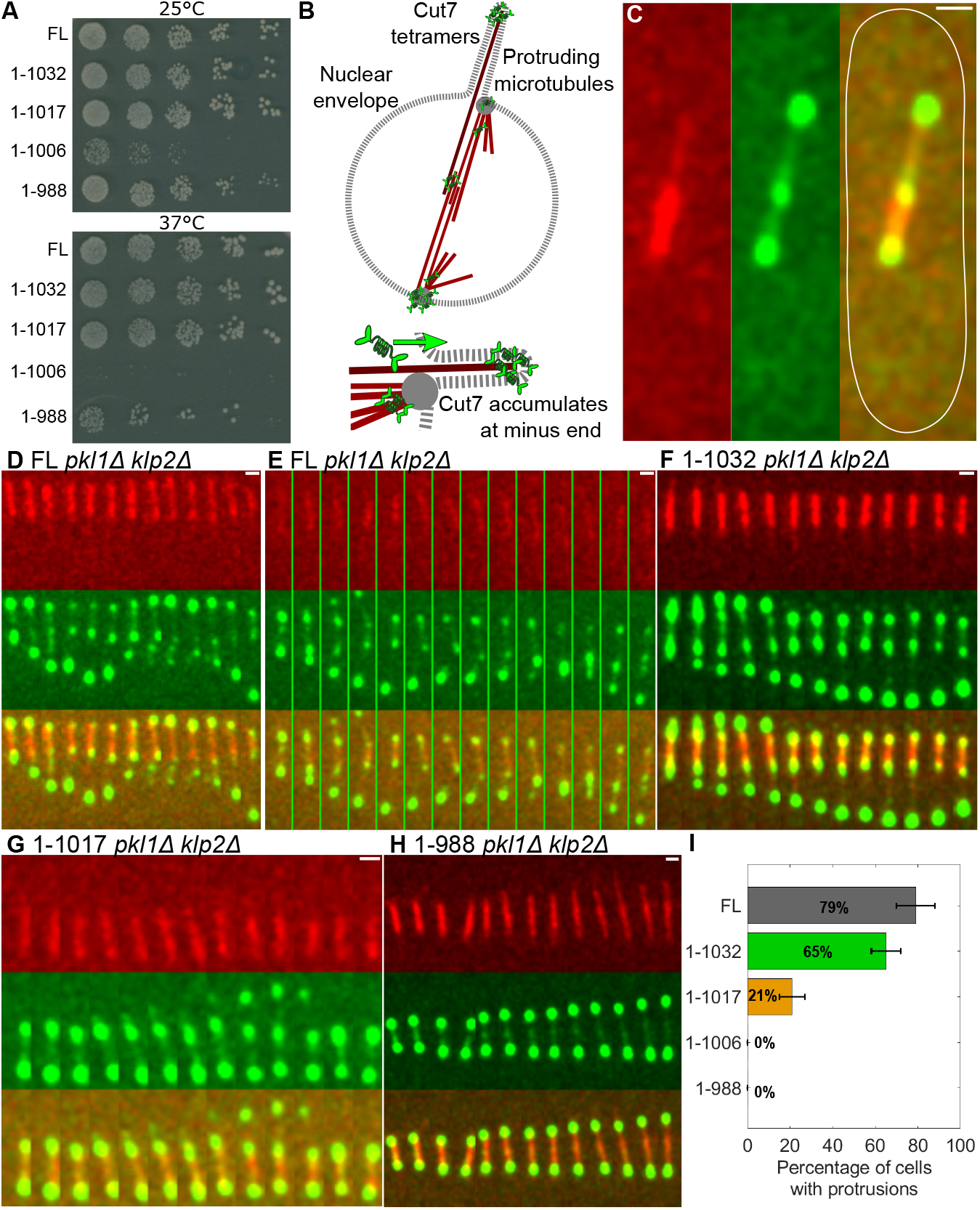
Kinesin-14 deletion rescued lethal *cut7* tail truncation mutants and led to spindle MT minus-end protrusions. (A) Serial dilution assay of truncation mutants in the *pkl1*Δ *klp2*Δ background 25°C (upper) and 37°C (lower). Cells with *cut7-1006* and *cut7-988* were viable but showed temperature-sensitive growth inhibition or lethality. (B) Schematic of protrusions and Cut7-GFP accumulation. Upper, MT with its minus-end detached from the SPB protrudes beyond the SPB and distorts the nuclear envelope. (Lower), Cut7-GFP near SPBs moves toward minus ends of protruding MT and collects at the protrusion tip. (C) Fluorescence image showing mCherry-atb2 MTs (red, left), Cut7-GFP (green, center), and merge (right) for cell grown and imaged at 25°C and shown as maximum-intensity projections of confocal sections. (D-H) Time-lapse imaging of spindle and protrusion dynamics showing mCherry-atb2 MTs (red, upper), Cut7-GFP (green, center), and merge (lower) for cells grown and imaged at 25°C and shown as maximum-intensity projections of confocal sections. (D-F) Protrusions in *cut7-FL* and *cut7-1032* cells. (G) Protrusion in *cut7-1017* cell. (H) Spindle in *cut7-988* cell showing no protrusions. Scalebar, 1 *µ*m. Time between images 82.4s (D), 82.4s (E), 27.4s (F), 36.8s (G), and 35.8s (H). (I) Fraction of cells with visible protrusions decreased as more of the tail is truncated. Data from 75 *cut7*, 43 *cut7-1032*, 39 *cut7-1017*, and 39 *cut7-1006* and 39 *cut7-988* cells. Error bars were estimated from Poisson counting statistics.

### Dynamic spindle MT minus-end protrusions serve as a read-out of net sliding force produced by Cut7 mutants

Cut7p outward-directed force generated at the midzone of the spindle is required for protrusion of spindle MT minus ends in fission yeast lacking kinesin-14/Pkl1 (or members of its complex, Wdr8p and Msd1p) [52–54, 61]. This suggested that protrusions might vary for Cut7p truncation mutants that caused changes in spindle length. Therefore, we imaged protrusions for our truncation mutants in the *pkl1*Δ *klp2*Δ background (Fig. 4B-H).

We found that Cut7p produced protrusions and accumulated at protrusion tips. We observed protrusions at 25°C in cells with fluorescently labeled MTs and Cut7p, as seen previously [52, 53]. In cells with protrusions, Cut7-GFP localized both to the SPBs and at the tip of the protruding MTs (Fig. 4B, C). Time-lapse imaging showed that some protrusions grew, shortened, and then grew again in the same cell (Fig. 4D), while other cells showed protrusions from both SPBs simultaneously (Fig. 4E). Because we used low-level tubulin labeling [58, 69] and protrusions contain only a small number of MTs [52], protrusion MTs were often too dim to be be visible in every image (Fig. 4D-H). Despite this, Cut7-GFP at protrusion tips was consistently bright, allowing identification of protrusions and their dynamics (Fig. 4C-H). Because protruding MTs typically have their minus-ends distal to the nucleus [52, 54], Cut7-GFP accumulation at protrusion tips appeared to arise from Cut7p localized at MT minus ends (Fig. 4B, lower). We noted that the protrusion tip often showed brighter GFP signal than the adjacent SPB (Fig. 4D-F).

Cut7p tail truncations showed reduced frequency of protrusions, even for alleles with mild growth and spindle phenotypes. In the *pkl1+ klp2+* background, *cut7-1032* and *cut7-1017* alleles with shorter tail truncation showed growth, spindle length, and midzone localization similar to *cut7-FL* (Fig. 2, 3). These observations suggested that the mild truncations could generate force similarly to full-length protein. However, when we measured the fraction of cells with visible protrusions, we found a decrease in all truncation mutants. Cells containing *cut7-FL* showed protrusions in 78% of cells, but this dropped to 65% for *cut7-1032* and 21% for *cut7-1017* (Fig. 4F,G,I). The removal of the residues C-terminal to the BimC motif in *cut7-1017* therefore appeared to cause a sharp decrease in net Cut7p sliding force. When the BimC motif was completely removed in *cut7-1006* and *cut7-988*, we did not observe any protrusions (Fig. 4H,I). Thus, protrusion frequency was strongly correlated with the amount of C-terminal tail present.

### Different portions of the Cut7p tail affect sliding force and midzone localization

We asked whether the changes in protrusion frequency in *cut7* trail truncations might be correlated with changes in spindle length or Cut7p localization. Therefore, we analyzed spindle length dynamics and Cut7-GFP intensity distribution for truncation mutants in the *pkl1*Δ *klp2*Δ background (Methods). The spindle sometimes shortened when protrusions formed, suggesting that formation of protrusions altered spindle force balance. Therefore, when measuring spindle length dynamics, we only analyzed cells without visible protrusions. Among these cells, pre-anaphase spindle length was shorter in *cut7-1006* and *cut7-988* cells lacking the BimC motif (Figs. 5A, S8). Since these strains also did not show protrusions in cells we imaged (Fig. 4I), this result could occur if both shorter spindle length and loss of protrusions result from lower net Cut7p sliding force.

**FIG. 5.**
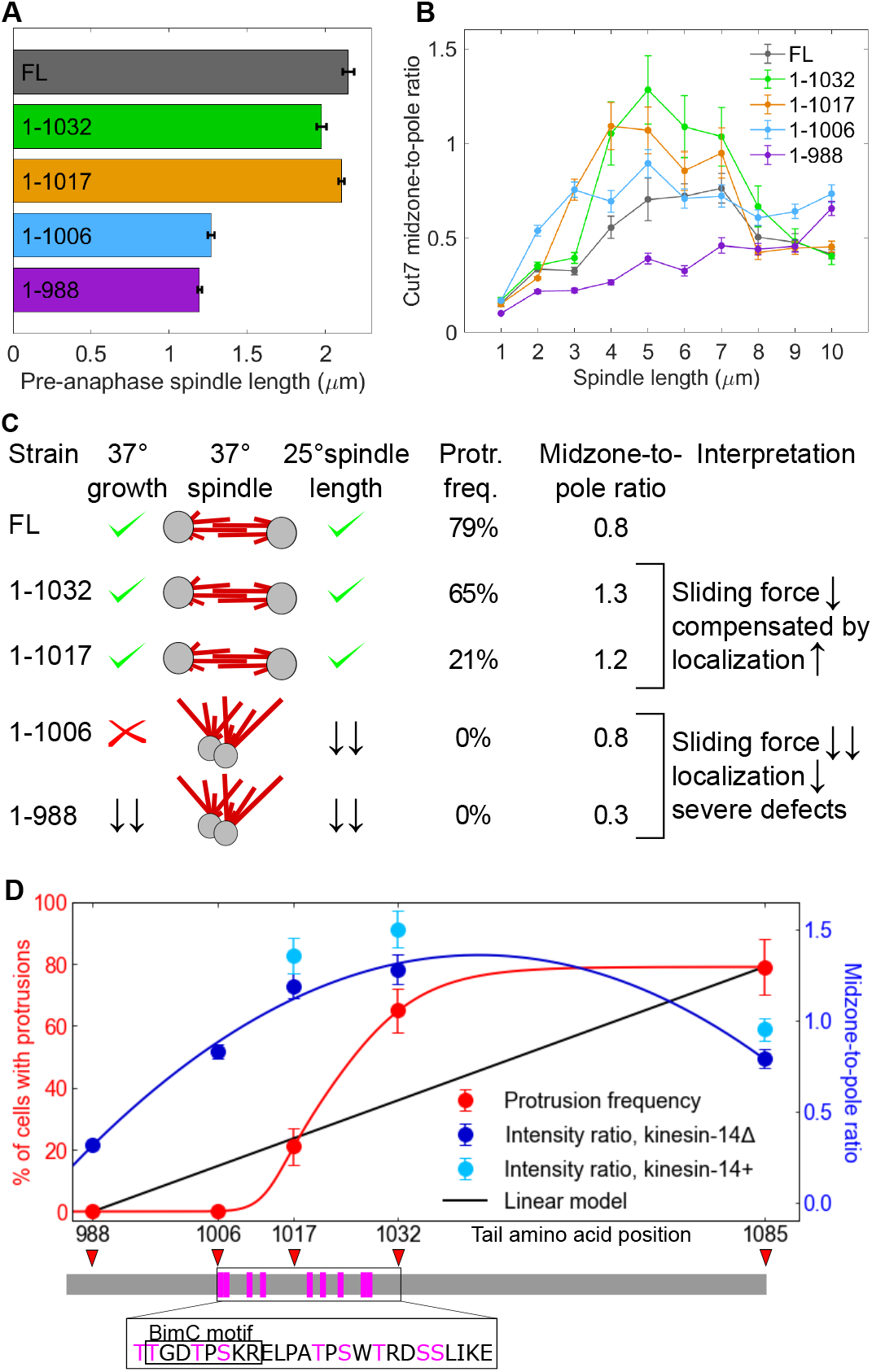
Spindle length, Cut7-GFP midzone-to-pole intensity ratio, and protrusion frequency varied non-linearly with tail truncation in the *pkl1*Δ *klp2*Δ background. (A) Pre-anaphase spindle length. Data from 14 *cut7*, 13 *cut7-1032*, 11 *cut7-1017*, 6 *cut7-1006*, and 17 *cut7-988* cells that did not exhibit protrusions (Methods). Error bars are standard error of the mean. (B) Cut7-GFP midzone-to-pole ratio as a function of spindle length. Intensity data from 44 *cut7*, 34 *cut7-1032*, 35 *cut7-1017*, 13 *cut7-1006*, and 40 *cut7-988* cells. (C) Summary of phenotypes of tail truncation mutants in the *pkl1*Δ *klp2*Δ deletion background, showing for each strain growth and spindle phenotypes at 37°C, pre-anaphase spindle length at 25°C, protrusion frequency, and average midzone-to-pole ratio. (D) Protrusion frequency (red) and midzone-to-pole intensity ratio (dark blue, *pkl1*Δ *klp2*Δ background; light blue, *pkl1+ klp2+* background) as a function of position of tail truncation. Black line, linear model. Error bars of protrusion frequency were estimated from Poisson counting statistics, error bars of intensity ratio were computed as the standard error of the mean assuming independent intensity measurements for each images at each integer spindle length. Lower, Cut7 tail with truncation sites marked with red triangles and phosphorylation sites with pink boxes. Inset: sequence and phosphorylated residues for a.a. 1006-1032.

The net sliding force could decrease in tail truncation mutants if the loss of tail amino acids lowers motor binding affinity for spindle microtubules, even if the net force per motor is unchanged in the mutants. We therefore asked whether the apparent reduction in net force might be caused by lower overall Cut7p levels or a reduced amount of motor at the spindle midzone. For this analysis we quantified all cells, with and without protrusions. Because protrusions complicated automated identification of spindle poles, we used manual tracking of SPB position in cells with visible protrusions (Methods). The total Cut7-GFP signal on the entire spindle was similar for all alleles, except for Cut7-988-GFP which showed *higher* total intensity (Fig. S9A). The Cut7-GFP midzone-to-pole ratio for most truncation alleles was similar to or higher than Cut7-FL, with the exception of Cut7-988-GFP which showed a consistently lower ratio than the other strains (Fig. 5B, S9B-F). Therefore, Cut7-988-GFP showed enhanced pole localization and decreased midzone localization, in agreement with previous observations [34].

Our analysis showed that Cut7-GFP midzone localization was increased or retained in most truncation mutants, despite the decrease in protrusions. To quantify localization, we averaged the Cut7-GFP midzone-to-pole ratio (Fig. 5B) for all spindles with length from 3 to 7 *µ*m. This gave a single average midzone-to-pole ratio that we compared to other truncation phenotypes (Fig. 5C). This comparison showed that our mutants fell into two groups. The moderate truncations to a.a. 1032 or 1017 led to a decrease in protrusions but an increase in midzone-to-pole ratio, suggesting that any decrease in sliding force in these alleles (as measured by protrusions) may be compensated by greater relative amounts of motor in the midzone, leading to overall growth and spindle length similar to *cut7-FL* (Fig. 5C). On the other hand, the increased truncation to a.a. 1006 or 988 eliminated protrusions but had a similar or lower midzone-to-pole ratio compared to full length Cut7p, suggesting that these alleles had defects in sliding force that were not compensated by increased midzone localization that caused temperature-sensitive growth defects and monopolar spindles (Fig. 5C).

In addition, we observed that neither protrusion frequency nor midzone-to-pole ratio changed linearly with the length of C-terminal tail retained. Therefore, we compared their variation with that of a linear model (Fig. 5D). Midzone localization was increased or retained (average ratio 0.8 or higher) for all truncations except that at a.a. 988, even for the truncation at a.a. 1006 that removed the BimC motif. We noted that the midzone-to-pole ratio was similar for *cut7-1006* and *cut7-FL* cells in the *pkl1*Δ *klp2*Δ background. The BimC motif was required for protrusions to occur in the context of truncation mutants, and protrusion frequency increased sharply as the truncation site increased from a.a. 1006 to 1032. Therefore, our results show that among truncation mutants, a significant protrusion frequency requires both the BimC motif and adjacent C-terminal amino acids. This region includes nine residues previously identified as phosphorylated during mitosis in *S. pombe* [70–73]. These results suggest that a particularly important tail region for force production is a.a. 1006-1032 (the BimC motif and adjacent C-terminal residues), while for anaphase midzone localization a.a. 988-1006 (N-terminal to the BimC motif) are most important.

## DISCUSSION

Our results show that the fission-yeast kinesin-5/Cut7 C-terminal tail contributes to spindle midzone localization, spindle assembly, and spindle length. Sequential truncation of the tail caused growth defects, temperature-sensitive lethality due to a failure of spindle assembly, and lethality as more of the tail was truncated. In viable mutants, we observed that pre-anaphase spindle length decreased and the time to reach steady anaphase elongation increased, suggesting that partial tail truncation limits net Cut7p sliding force. This occurred despite spindle localization that increased at the midzone in these partial truncations, suggesting that increased midzone localization could partially rescue decreased sliding force per motor.

We built on the observation that fission yeast lacking kinesin-14/Pkl1 form spindle MT minus-end protrusions that distort the nuclear envelope [52–54]. We found that as more of the C-terminal tail was truncated, the frequency of visible protrusions decreased. Protrusion formation is driven by Cut7 sliding force and resisted by drag forces from MT crosslinking in the midzone. Therefore, the decrease of protrusions in tail truncation mutants could in principle reflect higher drag [17] (perhaps due to a more rigor-like state) or decreased sliding force. Previous work has shown that tail truncation decreases the sliding force of human Eg5 *in vitro* [18], while in fission yeast, protrusions are not observed in *pkl1*Δ cells lacking *cut7* [54] or carrying a temperature-sensitive *cut7* loss of function [53]. Therefore, we believe that decreased protrusions likely arise from lower sliding force in tail truncation mutants rather than increased drag. Since spindle and midzone localization is retained or even increased in most tail truncation mutants, our results suggest that the net sliding force per motor decreases as the tail is truncated.

To connect protrusion formation more directly to motor force, we estimated the force required for nuclear envelope deformation to create a protrusion. Previous modeling work found that the characteristic force scale to create and extend a cylindrical membrane tube of radius *R* is *f* = 2*πR*(*κ/*(2*R*^2^) + *σ*), where *σ* is the surface tension of the nuclear envelope and *κ* is its bending rigidity [64, 74, 75]. The *S. pombe* nuclear envelope has an estimated *σ* = 0.013 pN/nm and *κ* = 40 pN nm [76, 77]. Electron microscopy of a bundle with 3 protruding MTs showed a bundle radius of ∼70 nm [52], while that of other nuclear MT bundles found a protrusion radius of 50 nm [76]. A protrusion containing a single MT and space for a 55-nm long motor would give an estimated minimum radius of 40 nm, suggesting 40-70 nm as a range of protrusion radii. The force required to create a 40-nm radius tube is *f* = 6.4 pN, while for a 70-nm radius tube it is *f* = 7.5 pN. Therefore we estimate that creating a protrusion requires sliding force on an MT bundle of 6-8 pN.

While sliding force has not previously been measured for Cut7p, it has been quantified in optical trapping measurements of antiparallel MT sliding of other kinesin-5 motors. Budding-yeast Cin8 can produce ∼1.5 pN of sliding force per motor, with a force that increases for multiple motors up to ∼10 pN [78]. If Cut7p produces force similarly to Cin8, a handful of motors would be sufficient to create a protrusion. Ensembles of purified human Eg5 produced 4 pN of force per *µ*m of MT overlap; this was reduced to 0.5 pN/*µ*m in a C-terminal tail truncation mutant [18]. Our observation of decreased or absent protrusions in cells with Cut7p tail truncation could be explained by a reduction in Cut7p sliding force below ∼ 6 pN, consistent with the force decrease observed for the Eg5 tail truncation. This suggests that C-terminal tail truncation could decrease kinesin-5 sliding force similarly for both *S. pombe* and human motors. Cut7p tail truncation in principle could cause an increase in drag force, decreasing the net force due to higher friction.

These results extend recent work on homologous kinesin-5 motors in reconstituted systems. Tail truncation alters ATPase activity, motility, and force of *Drosophila* and human kinesin-5s [18], showing the tail plays a role in tuning kinesin-5 force. C-terminal tails may bridge to adjacent motors, promoting cluster formation that could affect sliding force [50] via motor domain structural change upon tail binding [18]. The lethality of the *cut7* tail truncation we found is consistent with studies of kinesin-5/Cin8 in budding yeast [51]. A computational model of budding yeast Cin8 proposed an attractive interaction that promotes motor clustering [33], which could be mediated by tail-motor interactions.

Our data suggest that midzone localization and sliding force are affected by different portions of the Cut7p tail in truncation mutants. As has been seen with other *cut7* tail mutants [41], all of our truncated Cut7p alleles localized to spindle MTs/SPBs, despite the loss of the tail-MT binding site [43]. The localization, spindle length, and protrusion measurements together point to amino acids N-terminal to the BimC motif as most important for midzone localization (a.a. 988-1006), while the BimC motif and adjacent C-terminal amino acids are essential for sliding force (a.a. 1006-1032) in the context of tail truncation. The BimC motif (a.a. 1008-1015) contains a conserved Cdk phosphorylation site [5, 41, 42, 45, 46, 56]. When the Cdk-site threonine is mutated in vertebrate kinesin-5s, motor-spindle MT binding is reduced [5, 42, 46]. For Cut7p, spindle localization is retained with loss of the BimC motif, but net sliding force appears to be reduced. Nine sites phosphorylated in mitosis have been identified in the *cut7* tail between amino acids 1007 and 1028 [70–73]. This corresponds remarkably well with the region most important for protrusions in truncation mutants. We therefore speculate that phosphoregulation at these sites contributes to Cut7p sliding force.

While we did not see significant phenotypes of truncating amino acids 1033-1085 C-terminal to the BimC box region, these residues are still significant in the context of the internal deletion. The *cut7-988* and *cut7-989-1028*Δ alleles both lack the BimC motif, but only the full tail truncation shows a lethal phenotype. This suggests that even with this important region removed, retaining the other 57 tail amino acids is beneficial. Similarly, the internal deletion shows higher midzone-to-pole ratio compared to the full tail truncation, suggesting that the presence of the entire tail contributes to Cut7p midzone localization.

Tail truncation affected pre-anaphase spindle length and protrusion frequency, consistent with alterations in Cut7p force early in mitosis. While the truncation mutants did cause delays in reaching steady anaphase elongation, the late-anaphase elongation speed was similar for the different mutants. This is consistent with previous work showing that kinesin-6/Klp9 is the primary spindle elongation motor in fission-yeast anaphase B [79, 80].

Our results give further evidence of Cut7p bidirectional motility in cells. Previous work found that Cut7p moves toward plus and minus ends of MTs both when purified [28–30] and on the spindle [34]. In reconstituted antiparallel MT overlaps, crowding on the MT lattice shifted Cut7p to plus-end-directed movement [30]. However, we observed most Cut7p localized near the SPBs for all of our mutants, consistent with minus-end-directed movement that concentrates the motor near spindle poles. At first glance, this may appear to be at odds with the idea that crowding regulates Cut7p plus-end-directed motility on spindle MTs. However, several factors may explain the difference in results. The limited resolution of our images does not rule out the possibility that motor crowding near the SPB may cause plus-end-directed movement where Cut7p concentration is highest, while faster minus-end-directed motility could still concentrate Cut7p near the poles. Tail-mediated interactions between Cut7p tetramers could also cause a local increase in concentration near the poles of motors not bound to spindle MTs.

## METHODS

### Reagents and techniques

Oligonucleotide primers were purchased from Integrated DNA Technologies (Coralville, IA). Restriction enzymes and Phusion HF DNA polymerase were purchased from New England Biolabs (Ipswich, MA). DNA was prepared using Qiaprep Spin Miniprep Kit and polymerase chain reaction (PCR) products were purified using Qiaquick PCR Purification Kit, both from Qiagen (Germantown, MD). Fission yeast genomic DNA was prepared using YeaStar Genomic DNA Kit from Zymo Research (Irvine, CA). DNA sequencing was performed by Quintarabio (Hayward, CA). DNA concentration was determined using a Thermo Scientific Nanodrop 2000. Strains were verified by PCR and sequence analysis of genomic DNA. At least two different transformant strains were analyzed in experiments, and cells were cultured using standard techniques [81]. Growth phenotypes were analyzed after 3-4 days on rich media plates with 5-fold serial dilution starting with 5 microliters of culture at 0.1 optical density.

### Strain construction for truncation mutants

The plasmid pFA6a-GFP(S65T)-kanmx6 (Addgene 39292) was amplified by PCR using oligonucleotide primers Cut7-Cterm-pFA6a-GFP-F and Cut7-Cterm-pFA6a-GFP-R. The truncation mutants were made using the same reverse primer as above and the following forward primers: for 1-1032: Cut7-1-1032-GFP-F, for 1-1017: Cut7-1-1017-GFP-F for 1-1006: Cut7-1-1007-GFP-F, for 1-988: Cut7-1-988-GFP-F. There is a six amino acid linker (RIPGLI) between Cut7 and GFP. The PCR products were used for transformation using a lithium acetate method [82] and G418 resistant colonies were characterized. Transformation of yMB1084 (containing *pkl1+ klp2+*) yielded strains yMB1149 (*cut7*), yMB1125 (*cut7-1032*, and yMB1142 (*cut7-1017*) and transformation of strain yMB1147 (containing *pkl1*Δ *klp2*Δ) yielded strains yMB1162 (*cut7*), yMB1210 (*cut7-1032*), yMB1207 (*cut7-1017*), yMB1238 (*cut7-1006*) and yMB1155 (*cut7-988*).

### Strain construction for internal deletion mutant

Plasmid pMB64 was made in a Bluescript backbone (kind gift from Wayne Wahls) and contains the *cut7* coding sequence from Bgl2 (b.p. 1835) to the C-terminus (b.p. 3255) and downstream sequence through the *his1* gene. Synthetic *cut7* alleles (Genscript) were inserted into pMB64, from Bcl1 (b.p. 2356) to BsrG1 (b.p. 3361) containing either wild-type sequence with four silent restriction sites (pMB71-FL) or a *cut7* fragment containing an internal deletion, lacking amino acids 989-1028. These were cloned into a related plasmid containing a C-terminal GFP (pMB44-FL and pMB45-intdel respectively) and a fragment from these plasmids was transformed into yeast strain MB1091 (in which the downstream his1 gene had been replaced with ura4) and *his+ ura-*colonies were characterized. This yielded strains yMB1199 (*cut7*) and yMB1168 (*cut7-989-1028*Δ). Yeast with a *cut7* allele containing silent restriction sites (yMB1199) grew the same and behaved similarly in all assays as those with an allele lacking silent restriction sites (yMB1149). Therefore yMB1149 was used as the FL control for experiments in kinesin-14+ background.

### Live-cell imaging

All microscopy used live-cell preparation on a Nikon Eclipse Ti (Nikon USA, Melville, NY) spinning-disk confocal microscope as previously described [34, 64]. Microscope temperature was maintained with *±*0.1°C precision using a CherryTemp controller (Cherry Biotech, Rennes, France), and samples were placed on *Bandeiraea simplicifolia* lectin (Sigma-Aldrich, St. Louis, MO) coated coverslips pre-equilibrated to the appropriate temperature of either 25 or 37°C as described previously [34].

### Kymograph generation

For each truncation mutant, we selected the mitotic cells that formed a bipolar spindle, based on the bright spots of Cut7-GFP localized at both spindle poles. We manually segmented an elliptical region to include our cell of interest; any region outside this segmentation was not considered. The position of both Cut7-GFP spots was tracked to sub-pixel accuracy for each mitotic frame. We constructed kymographs by interpolating the Cut7-GFP intensity along the line connecting the spots in each frame and subtracting the background intensity. The brighter spot was used as a fixed reference point plotted on the left side of the kymograph, while the other spot varied in position. In some cases the line along the spindle axis extends beyond the boundary of the cell, into the region that has been cropped. In this case the kymograph includes pixels that have been cropped and therefore have zero intensity, which are visible in the kymographs as completely black areas.

### Cut7-GFP intensity analysis

To group profiles of Cut7-GFP intensity along the spindle, we identified the Cut7-GFP peaks near the SPBs and measured the distance between the peaks. We then extracted lines from the kymographs with the relevant peak separation. We included images with peaks separated by *±*0.1 *µ*m around the desired peak separation. In average curves, the right peak typically showed lower intensity than the left peak. This occurred because we manually selected the brighter Cut7-GFP spot to be at *x* = 0. We computed total Cut7-GFP intensity by summing the intensity from each spindle intensity profile extracted from the kymographs.

We determined the Cut7-GFP midzone-to-pole ratio by first defining the pole and midzone. We summed pole intensity from a square box 0.75 *µ*m on a side centered on the pole intensity peak. The midzone intensity is summed in a rectangular box of width 0.75 *µ*m with long axis centered on the spindle axis that runs between the two poles, including all of the spindle except the pole boxes. The total Cut7-GFP intensity is the sum of the pole and midzone signals. The midzone-to-pole ratio is the total midzone Cut7-GFP intensity, divided by total pole intensity, computed for each frame.

### Spindle length dynamics

Microscope images were processed using TAMiT [60], which identifies spindle endpoints in each frame. The length of the spindle is determined from these points and smoothed. First, we divide each trace into 30 time bins and analyzed the standard deviation of spindle length in each bin. Bins with high standard deviation (corresponding to fluctuations on the order of 0.75 *µ*m) were flagged as noise and discarded. The data were then smoothed with a rolling average with window of width ∼3.5 min.

We manually identified anaphase onset for each curve by choosing the point where the spindle begins to grow beyond fluctuations around its pre-anaphase length. To produce average spindle length traces from different cells, each curve was aligned by its anaphase onset time and divided into evenly spaced bins of width 15 s. All points from each curve that fall within a given bin were then averaged. Average curves for each phenotype included at least 4 curves.

To determine the anaphase elongation rate, we first manually identified the time range of constant growth speed, then performed a linear fit on this region. The fit elongation speed was averaged from multiple cells. The average time of onset of steady anaphase elongation was determined by averaging the identified time from all cells. Uncertainty is computed and displayed as the standard error of the mean.

## ACKNOWLEDGMENTS

We thank Daniel Steckhahn for helpful discussions and figure formatting, the University of Colorado Boulder Light Microscopy and Biocore Facilities, Pombase and the Wellcome Trust, and the National Bio-Resource Project Yeast, Japan for providing original strains. This work was funded by NIH grant R01GM124371 and NSF grant MCB 2133243.

## References

[1] A. P. Enos and N. R. Morris, Mutation of a gene that encodes a kinesin-like protein blocks nuclear division in A. nidulans, Cell 60, 1019 (1990).

[2] I. Hagan and M. Yanagida, Novel potential mitotic motor protein encoded by the fission yeast cut7+ gene, Nature 347, 563 (1990).

[3] M. A. Hoyt, L. He, K. K. Loo, and W. S. Saunders, Two Saccharomyces cerevisiae kinesin-related gene products required for mitotic spindle assembly., The Journal of Cell Biology 118, 109 (1992).

[4] K. E. Sawin, K. LeGuellec, M. Philippe, and T. J. Mitchison, Mitotic spindle organization by a plus-end-directed microtubule motor, Nature 359, 540 (1992).

[5] A. Blangy, H. A. Lane, P. d’Hérin, M. Harper, M. Kress, and E. A. Nigg, Phosphorylation by p34cdc2 regulates spindle association of human Eg5, a kinesin-related motor essential for bipolar spindle formation in vivo, Cell 83, 1159 (1995).

[6] D. G. Cole, W. M. Saxton, K. B. Sheehan, and J. M. Scholey, A “slow” homotetrameric kinesin-related motor protein purified from Drosophila embryos., Journal of Biological Chemistry 269, 22913 (1994).

[7] A. S. Kashina, R. J. Baskin, D. G. Cole, K. P. Wedaman, W. M. Saxton, and J. M. Scholey, A bipolar kinesin, Nature 379, 270 (1996).

[8] D. M. Gordon and D. M. Roof, The Kinesin-related Protein Kip1p of Saccharomyces cerevisiae Is Bipolar, Journal of Biological Chemistry 274, 28779 (1999).

[9] S. Acar, D. B. Carlson, M. S. Budamagunta, V. Yarov-Yarovoy, J. J. Correia, M. R. Niñonuevo, W. Jia, L. Tao, J. A. Leary, J. C. Voss, J. E. Evans, and J. M. Scholey, The bipolar assembly domain of the mitotic motor kinesin-5, Nature Communications 4, ncomms2348 (2013).

[10] J. E. Scholey, S. Nithianantham, J. M. Scholey, and J. Al-Bassam, Structural basis for the assembly of the mitotic motor Kinesin-5 into bipolar tetramers, eLife 3, e02217 (2014).

[11] S. K. Singh, H. Pandey, J. Al-Bassam, and L. Gheber, Bidirectional motility of kinesin-5 motor proteins: Structural determinants, cumulative functions and physiological roles, Cellular and Molecular Life Sciences 75, 1757 (2018).

[12] D. J. Sharp, K. L. McDonald, H. M. Brown, H. J. Matthies, C. Walczak, R. D. Vale, T. J. Mitchison, and J. M. Scholey, The Bipolar Kinesin, KLP61F, Cross-links Microtubules within Interpolar Microtubule Bundles of Drosophila Embryonic Mitotic Spindles, The Journal of Cell Biology 144, 125 (1999).

[13] L. C. Kapitein, E. J. G. Peterman, B. H. Kwok, J. H. Kim, T. M. Kapoor, and C. F. Schmidt, The bipolar mitotic kinesin Eg5 moves on both microtubules that it crosslinks, Nature 435, 114 (2005).

[14] E. R. Hildebrandt, L. Gheber, T. Kingsbury, and M. A. Hoyt, Homotetrameric Form of Cin8p, a Saccharomyces cerevisiae Kinesin-5 Motor, Is Essential for Its in Vivo Function, Journal of Biological Chemistry 281, 26004 (2006).

[15] L. Tao, A. Mogilner, G. Civelekoglu-Scholey, R. Wollman, J. Evans, H. Stahlberg, and J. M. Scholey, A Homotetrameric Kinesin-5, KLP61F, Bundles Microtubules and Antagonizes Ncd in Motility Assays, Current Biology 16, 2293 (2006).

[16] S. M. J. L. van den Wildenberg, L. Tao, L. C. Kapitein, C. F. Schmidt, J. M. Scholey, and E. J. G. Peterman, The Homotetrameric Kinesin-5 KLP61F Preferentially Crosslinks Microtubules into Antiparallel Orientations, Current Biology 18, 1860 (2008).

[17] Y. Shimamoto, S. Forth, and T. M. Kapoor, Measuring Pushing and Braking Forces Generated by Ensembles of Kinesin-5 Crosslinking Two Microtubules, Developmental Cell 34, 669 (2015).

[18] T. Bodrug, E. M. Wilson-Kubalek, S. Nithianantham, A. F. Thompson, A. Alfieri, I. Gaska, J. Major, G. Debs, S. Inagaki, P. Gutierrez, L. Gheber, R. J. McKenney, C. V. Sindelar, R. Milligan, J. Stumpff, S. S. Rosenfeld, S. T. Forth, and J. Al-Bassam, The kinesin-5 tail domain directly modulates the mechanochemical cycle of the motor domain for anti-parallel microtubule sliding, eLife 9, e51131 (2020).

[19] G. Goshima and J. M. Scholey, Control of Mitotic Spindle Length, Annual Review of Cell and Developmental Biology 26, 21 (2010).

[20] B. J. Mann and P. Wadsworth, Kinesin-5 Regulation and Function in Mitosis, Trends in Cell Biology (2018).

[21] J. M. Scholey, G. Civelekoglu-Scholey, and I. Brust-Mascher, Anaphase B, Biology 5, 51 (2016).

[22] T. U. Mayer, T. M. Kapoor, S. J. Haggarty, R. W. King, S. L. Schreiber, and T. J. Mitchison, Small Molecule Inhibitor of Mitotic Spindle Bipolarity Identified in a Phenotype-Based Screen, Science 286, 971 (1999).

[23] J. Roostalu, C. Hentrich, P. Bieling, I. A. Telley, E. Schiebel, and T. Surrey, Directional Switching of the Kinesin Cin8 Through Motor Coupling, Science 332, 94 (2011).

[24] A. Gerson-Gurwitz, C. Thiede, N. Movshovich, V. Fridman, M. Podolskaya, T. Danieli, S. Lakämper, D. R. Klopfenstein, C. F. Schmidt, and L. Gheber, Directionality of individual kinesin-5 Cin8 motors is modulated by loop 8, ionic strength and microtubule geometry, The EMBO Journal 30, 4942 (2011).

[25] R. Avunie-Masala, N. Movshovich, Y. Nissenkorn, A. Gerson-Gurwitz, V. Fridman, M. Kõoivomägi, M. Loog, M. A. Hoyt, A. Zaritsky, and L. Gheber, Phospho-regulation of kinesin-5 during anaphase spindle elongation, Journal of Cell Science 124, 873 (2011).

[26] C. Thiede, V. Fridman, A. Gerson-Gurwitz, L. Gheber, and C. F. Schmidt, Regulation of bi-directional movement of single kinesin-5 Cin8 molecules, BioArchitecture 2, 70 (2012).

[27] V. Fridman, A. Gerson-Gurwitz, O. Shapira, N. Movshovich, S. Lakämper, C. F. Schmidt, and L. Gheber, Kinesin-5 Kip1 is a bi-directional motor that stabilizes microtubules and tracks their plus-ends in vivo, J Cell Sci 126, 4147 (2013).

[28] M. Edamatsu, Bidirectional motility of the fission yeast kinesin-5, Cut7, Biochemical and Biophysical Research Communications 446, 231 (2014).

[29] M. Edamatsu, Molecular properties of the N-terminal extension of the fission yeast kinesin-5, Cut7., Genetics and molecular research : GMR 15 (2016).

[30] M. Britto, A. Goulet, S. Rizvi, O. von Loeffelholz, C. A. Moores, and R. A. Cross, Schizosaccharomyces pombe kinesin-5 switches direction using a steric blocking mechanism, Proceedings of the National Academy of Sciences, 201611581 (2016).

[31] O. Shapira and L. Gheber, Motile properties of the bi-directional kinesin-5 Cin8 are affected by phosphorylation in its motor domain, Scientific Reports 6, 25597 (2016).

[32] O. Shapira, A. Goldstein, J. Al-Bassam, and L. Gheber, Possible physiological role for bi-directional motility and motor clustering of the mitotic kinesin-5 Cin8, J Cell Sci, jcs.195040 (2017).

[33] H. Pandey, E. Reithmann, A. Goldstein-Levitin, J. Al-Bassam, E. Frey, and L. Gheber, Drag-induced directionality switching of kinesin-5 Cin8 revealed by cluster-motility analysis, Science Advances 7, eabc1687 (2021).

[34] Z. R. Gergely, S. Ansari, M. H. Jones, B. Zhou, C. Cash, R. McIntosh, and M. D. Betterton, Kinesin-5/Cut7 moves bidirectionally on fission-yeast spindles with activity that increases in anaphase, Journal of Cell Science, jcs.260474 (2023).

[35] M. Uteng, C. Hentrich, K. Miura, P. Bieling, and T. Surrey, Poleward transport of Eg5 by dynein–dynactin in Xenopus laevis egg extract spindles, The Journal of Cell Biology 182, 715 (2008).

[36] N. Ma, U. S. Tulu, N. P. Ferenz, C. Fagerstrom, A. Wilde, and P. Wadsworth, Poleward Transport of TPX2 in the Mammalian Mitotic Spindle Requires Dynein, Eg5, and Microtubule Flux, Molecular Biology of the Cell 21, 979 (2010).

[37] N. Ma, J. Titus, A. Gable, J. L. Ross, and P. Wadsworth, TPX2 regulates the localization and activity of Eg5 in the mammalian mitotic spindle, J Cell Biol 195, 87 (2011).

[38] A. Gable, M. Qiu, J. Titus, S. Balchand, N. P. Ferenz, N. Ma, E. S. Collins, C. Fagerstrom, J. L. Ross, G. Yang, and P. Wadsworth, Dynamic reorganization of Eg5 in the mammalian spindle throughout mitosis requires dynein and TPX2, Molecular Biology of the Cell 23, 1254 (2012).

[39] S. K. Balchand, B. J. Mann, J. Titus, J. L. Ross, and P. Wadsworth, TPX2 Inhibits Eg5 by Interactions with Both Motor and Microtubule, Journal of Biological Chemistry 290, 17367 (2015).

[40] I. Hagan and M. Yanagida, Kinesin-related cut 7 protein associates with mitotic and meiotic spindles in fission yeast, Nature 356, 74 (1992).

[41] D. R. Drummond and I. M. Hagan, Mutations in the bimC box of Cut7 indicate divergence of regulation within the bimC family of kinesin related proteins, Journal of Cell Science 111, 853 (1998).

[42] J. Cahu, A. Olichon, C. Hentrich, H. Schek, J. Drinjakovic, C. Zhang, A. Doherty-Kirby, G. Lajoie, and T. Surrey, Phosphorylation by Cdk1 Increases the Binding of Eg5 to Microtubules In Vitro and in Xenopus Egg Extract Spindles, PLOS ONE 3, e3936 (2008).

[43] Z. T. Olmsted, A. G. Colliver, T. D. Riehlman, and J. L. Paluh, Kinesin-14 and kinesin-5 antagonistically regulate microtubule nucleation by γ-TuRC in yeast and human cells, Nature Communications 5, 5339 (2014).

[44] R. Blackwell, C. Edelmaier, O. Sweezy-Schindler, A. Lamson, Z. R. Gergely, E. O’Toole, A. Crapo, L. E. Hough, J. R. McIntosh, M. A. Glaser, and M. D. Betterton, Physical determinants of bipolar mitotic spindle assembly and stability in fission yeast, Science Advances 3, e1601603 (2017).

[45] M. M. Heck, A. Pereira, P. Pesavento, Y. Yannoni, A. C. Spradling, and L. S. Goldstein, The kinesin-like protein KLP61F is essential for mitosis in Drosophila., The Journal of Cell Biology 123, 665 (1993).

[46] K. E. Sawin and T. J. Mitchison, Mutations in the kinesin-like protein Eg5 disrupting localization to the mitotic spindle, Proceedings of the National Academy of Sciences 92, 4289 (1995).

[47] J. Rapley, M. Nicolàs, A. Groen, L. Regué, M. T. Bertran, C. Caelles, J. Avruch, and J. Roig, The NIMA-family kinase Nek6 phosphorylates the kinesin Eg5 at a novel site necessary for mitotic spindle formation, Journal of Cell Science 121, 3912 (2008).

[48] E. Saeki, S. Yasuhira, M. Shibazaki, H. Tada, M. Doita, T. Masuda, and C. Maesawa, Involvement of C-terminal truncation mutation of kinesin-5 in resistance to kinesin-5 inhibitor, PLOS ONE 13, e0209296 (2018).

[49] J. S. Weinger, M. Qiu, G. Yang, and T. M. Kapoor, A nonmotor microtubule binding site in kinesin-5 is required for filament crosslinking and sliding, Current Biology 21, 154 (2011).

[50] S. Nithianantham, M. K. Iwanski, I. Gaska, H. Pandey, T. Bodrug, S. Inagaki, J. Major, G. J. Brouhard, L. Gheber, S. S. Rosenfeld, S. Forth, A. G. Hendricks, and J. Al-Bassam, The mechanochemical origins of the microtubule sliding motility within the kinesin-5 domain organization (2021).

[51] A. Düselder, V. Fridman, C. Thiede, A. Wiesbaum, A. Goldstein, D. R. Klopfenstein, O. Zaitseva, M. E. Janson, L. Gheber, and C. F. Schmidt, Deletion of the Tail Domain of the Kinesin-5 Cin8 Affects Its Directionality, Journal of Biological Chemistry 290, 16841 (2015).

[52] M. Toya, M. Sato, U. Haselmann, K. Asakawa, D. Brunner, C. Antony, and T. Toda, γ-Tubulin complex-mediated anchoring of spindle microtubules to spindle-pole bodies requires Msd1 in fission yeast, Nature Cell Biology 9, 646 (2007).

[53] M. Yukawa, C. Ikebe, and T. Toda, The Msd1–Wdr8–Pkl1 complex anchors microtubule minus ends to fission yeast spindle pole bodies, J Cell Biol 209, 549 (2015).

[54] V. Syrovatkina and P. T. Tran, Loss of kinesin-14 results in aneuploidy via kinesin-5-dependent microtubule protrusions leading to chromosome cut, Nature Communications 6, 7322 (2015).

[55] A. S. Rodriguez, A. N. Killilea, J. Batac, J. Filopei, D. Simeonov, I. Lin, and J. L. Paluh, Protein complexes at the microtubule organizing center regulate bipolar spindle assembly, Cell Cycle 7, 1246 (2008).

[56] T. Akera, Y. Goto, M. Sato, M. Yamamoto, and Y. Watanabe, Mad1 promotes chromosome congression by anchoring a kinesin motor to the kinetochore, Nature Cell Biology 17, 1124 (2015).

[57] O. von Loeffelholz, A. Pe ña, D. R. Drummond, R. Cross, and C. A. Moores, Cryo-EM Structure (4.5-Å) of Yeast Kinesin-5–Microtubule Complex Reveals a Distinct Binding Footprint and Mechanism of Drug Resistance, Journal of Molecular Biology 431, 864 (2019).

[58] H. A. Snaith, A. Anders, I. Samejima, and K. E. Sawin, Chapter 9 - New and Old Reagents for Fluorescent Protein Tagging of Microtubules in Fission Yeast: Experimental and Critical Evaluation, in Methods in Cell Biology, Vol. Volume 97, edited by Lynne Cassimeris and Phong Tran (Academic Press, 2010) pp. 147–172.

[59] R. Blackwell, O. Sweezy-Schindler, C. Baldwin, L. E. Hough, M. A. Glaser, and M. D. Betterton, Microscopic origins of anisotropic active stress in motor-driven nematic liquid crystals, Soft Matter 12, 2676 (2016).

[60] S. Ansari, Z. R. Gergely, P. Flynn, G. Li, J. K. Moore, and M. D. Betterton, Quantifying Yeast Microtubules and Spindles Using the Toolkit for Automated Microtubule Tracking (TAMiT), Biomolecules 13, 939 (2023).

[61] M. Yukawa, Y. Teratani, and T. Toda, How Essential Kinesin-5 Becomes Non-Essential in Fission Yeast: Force Balance and Microtubule Dynamics Matter, Cells 9, 1154 (2020).

[62] R. Ding, K. L. McDonald, and J. R. McIntosh, Three-dimensional reconstruction and analysis of mitotic spindles from the yeast, Schizosaccharomyces pombe., The Journal of Cell Biology 120, 141 (1993).

[63] J. J. Ward, H. Roque, C. Antony, and F. Nédélec, Mechanical design principles of a mitotic spindle, eLife 3, e03398 (2015).

[64] C. Edelmaier, A. R. Lamson, Z. R. Gergely, S. Ansari, R. Blackwell, J. R. McIntosh, M. A. Glaser, and M. D. Betterton, Mechanisms of chromosome biorientation and bipolar spindle assembly analyzed by computational modeling, eLife 9, e48787 (2020).

[65] A. L. Pidoux, M. LeDizet, and W. Z. Cande, Fission yeast pkl1 is a kinesin-related protein involved in mitotic spindle function., Molecular Biology of the Cell 7, 1639 (1996).

[66] S. A. Rincon, A. Lamson, R. Blackwell, V. Syrovatkina, V. Fraisier, A. Paoletti, M. D. Betterton, and P. T. Tran, Kinesin-5-independent mitotic spindle assembly requires the antiparallel microtubule crosslinker Ase1 in fission yeast, Nature Communications 8, 15286 (2017).

[67] M. Yukawa, Y. Yamada, T. Yamauchi, and T. Toda, Two spatially distinct Kinesin-14 Pkl1 and Klp2 generate collaborative inward forces against Kinesin-5 Cut7 in S. pombe, J Cell Sci, jcs.210740 (2018).

[68] A. R. Lamson, C. J. Edelmaier, M. A. Glaser, and M. D. Betterton, Theory of Cytoskeletal Reorganization during Cross-Linker-Mediated Mitotic Spindle Assembly, Biophysical Journal 116, 1719 (2019).

[69] R. Blackwell, O. Sweezy-Schindler, C. Edelmaier, Z. R. Gergely, P. J. Flynn, S. Montes, A. Crapo, A. Doostan, J. R. McIntosh, M. A. Glaser, and M. D. Betterton, Contributions of Microtubule Dynamic Instability and Rotational Diffusion to Kinetochore Capture, Biophysical Journal 112, 552 (2017).

[70] J. T. Wilson-Grady, J. Villén, and S. P. Gygi, Phosphoproteome Analysis of Fission Yeast, Journal of Proteome Research 7, 1088 (2008).

[71] A. Koch, K. Krug, S. Pengelley, B. Macek, and S. Hauf, Mitotic Substrates of the Kinase Aurora with Roles in Chromatin Regulation Identified Through Quantitative Phosphoproteomics of Fission Yeast, Science Signaling 4, rs6 (2011).

[72] A. N. Kettenbach, L. Deng, Y. Wu, S. Baldissard, M. E. Adamo, S. A. Gerber, and J. B. Moseley, Quantitative Phosphoproteomics Reveals Pathways for Coordination of Cell Growth and Division by the Conserved Fission Yeast Kinase Pom1, Molecular & Cellular Proteomics 14, 1275 (2015).

[73] M. P. Swaffer, A. W. Jones, H. R. Flynn, A. P. Snijders, and P. Nurse, Quantitative Phosphoproteomics Reveals the Signaling Dynamics of Cell-Cycle Kinases in the Fission Yeast Schizosaccharomyces pombe, Cell Reports 24, 503 (2018).

[74] I. Derényi, F. Jülicher, and J. Prost, Formation and Interaction of Membrane Tubes, Physical Review Letters 88, 238101 (2002).

[75] A. Lamson, C. Edelmaier, M. A. Glaser, and M. Betterton, Theory of cytoskeletal reorganization during crosslinker-mediated mitotic spindle assembly, bioRxiv, 419135 (2018).

[76] G. H. W. Lim, G. Huber, Y. Torii, A. Hirata, J. Miller, and S. Sazer, Vesicle-Like Biomechanics Governs Important Aspects of Nuclear Geometry in Fission Yeast, PLoS ONE 2, e948 (2007).

[77] G. H. W. Lim and G. Huber, The Tethered Infinitesimal Tori and Spheres Algorithm: A Versatile Calculator for Axisymmetric Problems in Equilibrium Membrane Mechanics, Biophysical Journal 96, 2064 (2009).

[78] T. Fallesen, J. Roostalu, C. Duellberg, G. Pruessner, and T. Surrey, Ensembles of Bidirectional Kinesin Cin8 Produce Additive Forces in Both Directions of Movement, Biophysical Journal 113, 2055 (2017).

[79] C. Fu, J. J. Ward, I. Loiodice, G. Velve-Casquillas, F. J. Nedelec, and P. T. Tran, Phospho-Regulated Interaction between Kinesin-6 Klp9p and Microtubule Bundler Ase1p Promotes Spindle Elongation, Developmental Cell 17, 257 (2009).

[80] M. Yukawa, M. Okazaki, Y. Teratani, K. Furuta, and T. Toda, Kinesin-6 Klp9 plays motor-dependent and -independent roles in collaboration with Kinesin-5 Cut7 and the microtubule crosslinker Ase1 in fission yeast, Scientific Reports 9, 7336 (2019).

[81] S. Moreno, A. Klar, and P. Nurse, Molecular genetic analysis of fission yeast Schizosaccharomyces pombe, in Guide to Yeast Genetics and Molecular Biology, Vol. Volume 194 (Academic Press, 1991) pp. 795–823.

[82] K. Okazaki, N. Okazaki, K. Kume, S. Jinno, K. Tanaka, and H. Okayama, High-frequency transformation method and library transducing vectors for cloning mammalian cDNAs by trans-complementation of Schizosaccharomyces pombe, Nucleic Acids Research 18, 6485 (1990).

