## Supplemental figures for "Distinct regions of the kinesin-5 C-terminal tail are essential for mitotic spindle midzone localization and sliding force"

#### This PDF file includes:

Supporting text
Figs. S1 to S9
Table S1

### Supporting Information Text

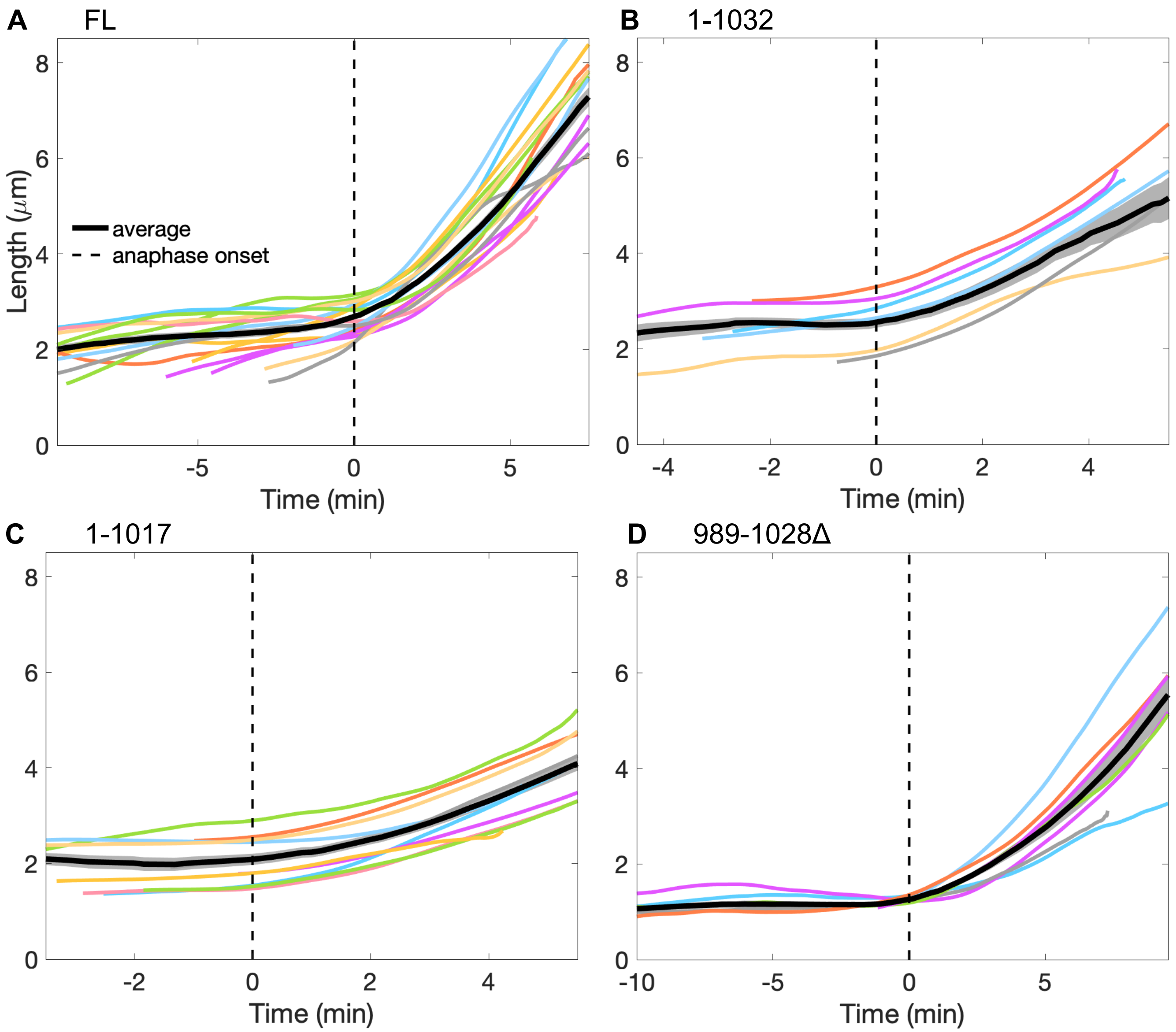


**Fig. S1.** Spindle length as a function of time showing curves from individual cells (colored lines) and average (black line). We defined $t = 0$ as the time of anaphase onset to align the curves (dashed line, Methods). Data from 19 *cut7* (A), 9 *cut7-1032* (B), 11 *cut7-1017* (C), and 14 *cut7-989-1028*Δ cells (D).

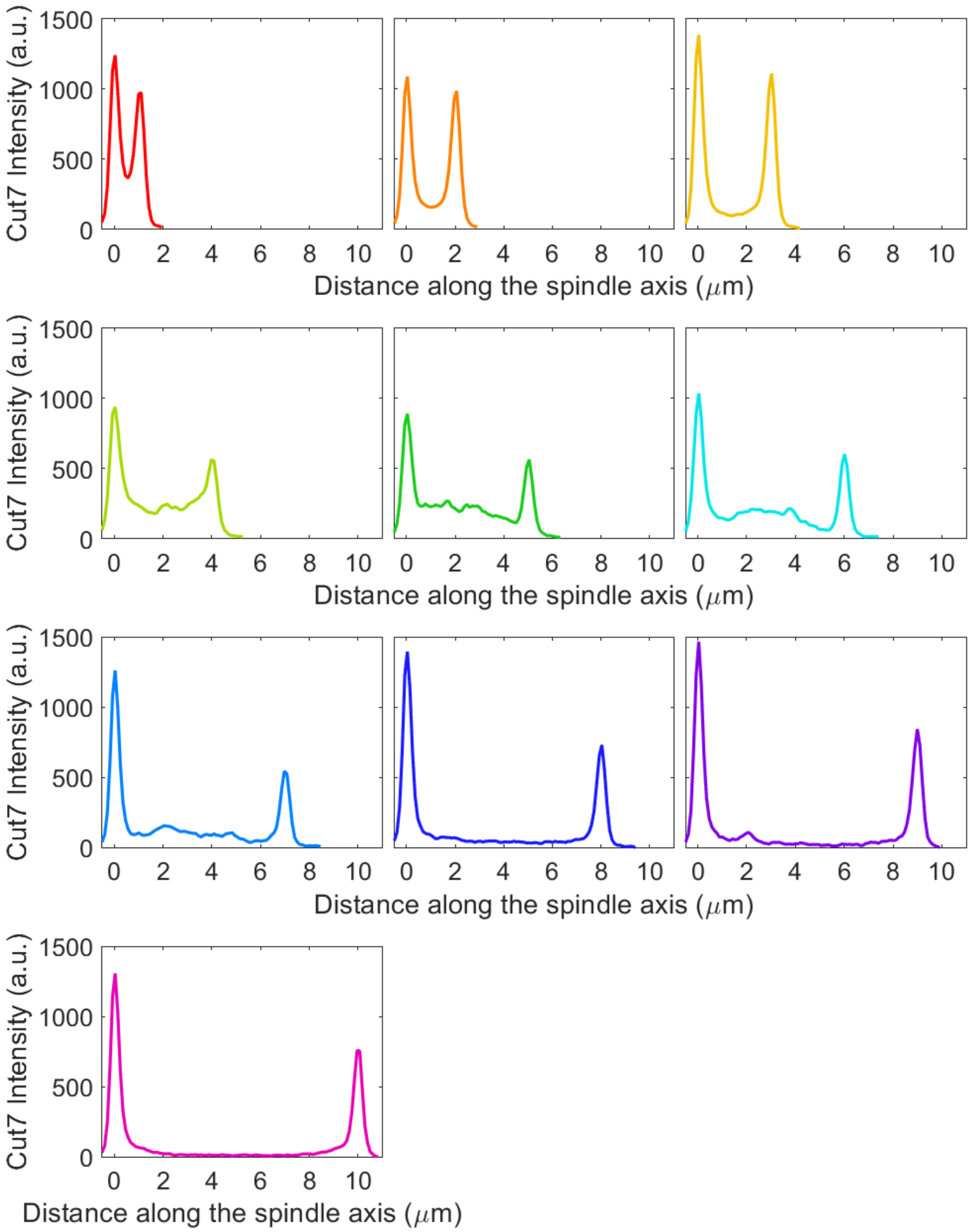


**Fig. S2.** Average Cut7-FL-GFP intensity along the spindle axis for varying spindle length. Intensity data from 29 *cut7-FL* cells.

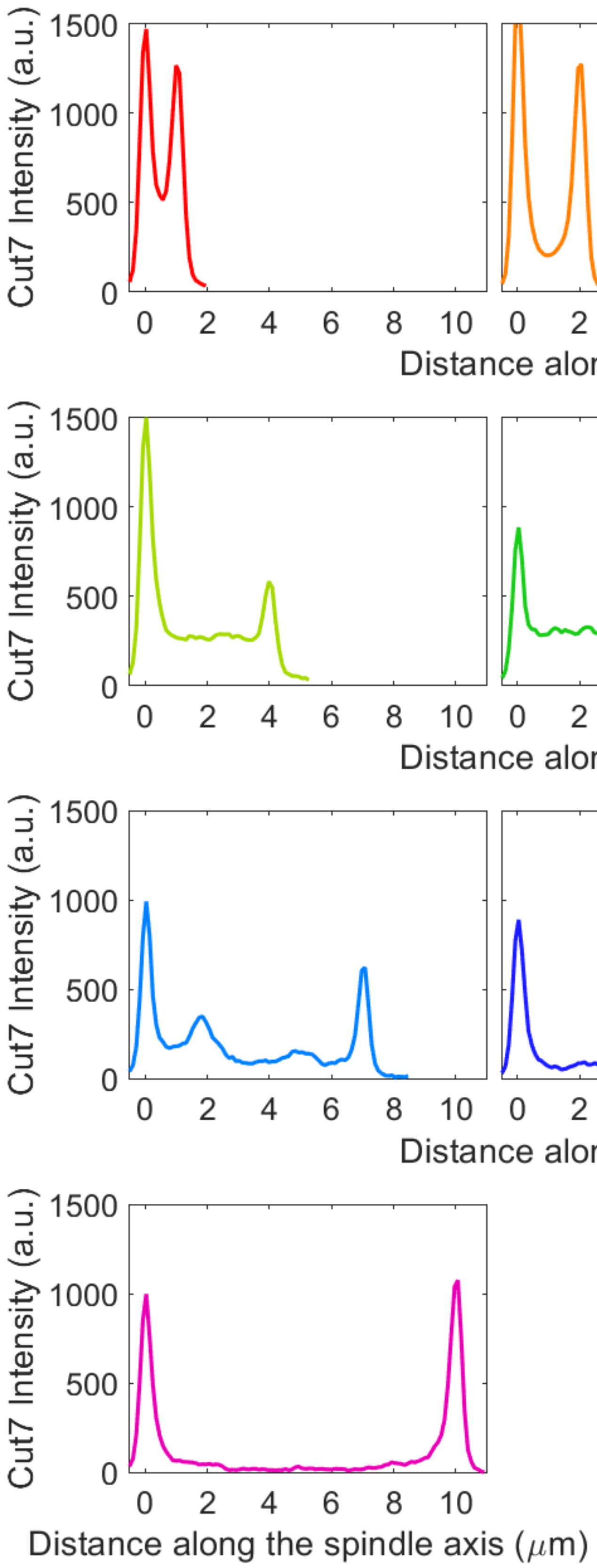


**Fig. S3.** Average Cut7-1032-GFP intensity along the spindle axis for varying spindle length. Intensity data from 34 *cut7-1032* cells.

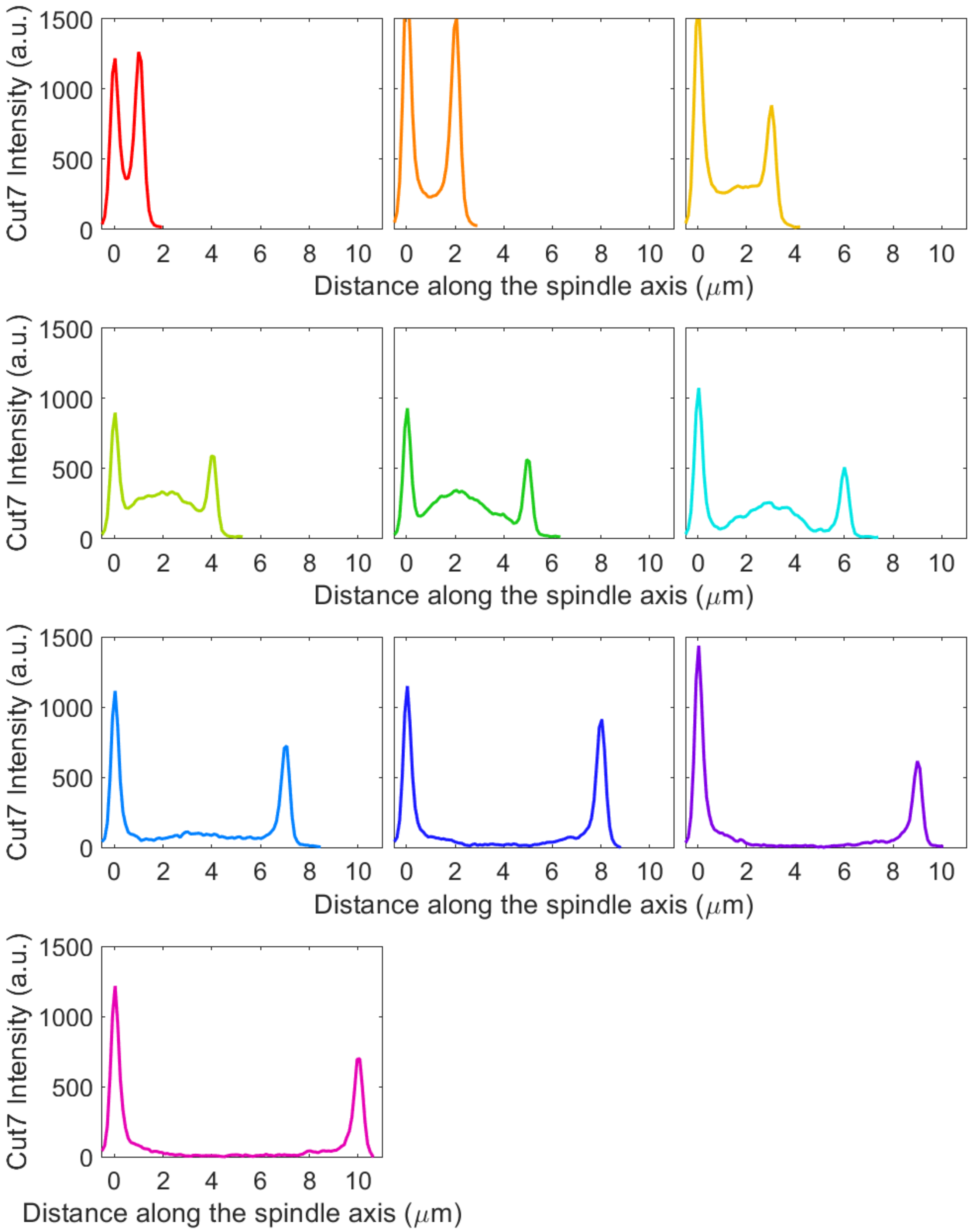


**Fig. S4.** Average Cut7-1017-GFP intensity along the spindle axis for varying spindle length. Intensity data from 29 *cut7-1017* cells.

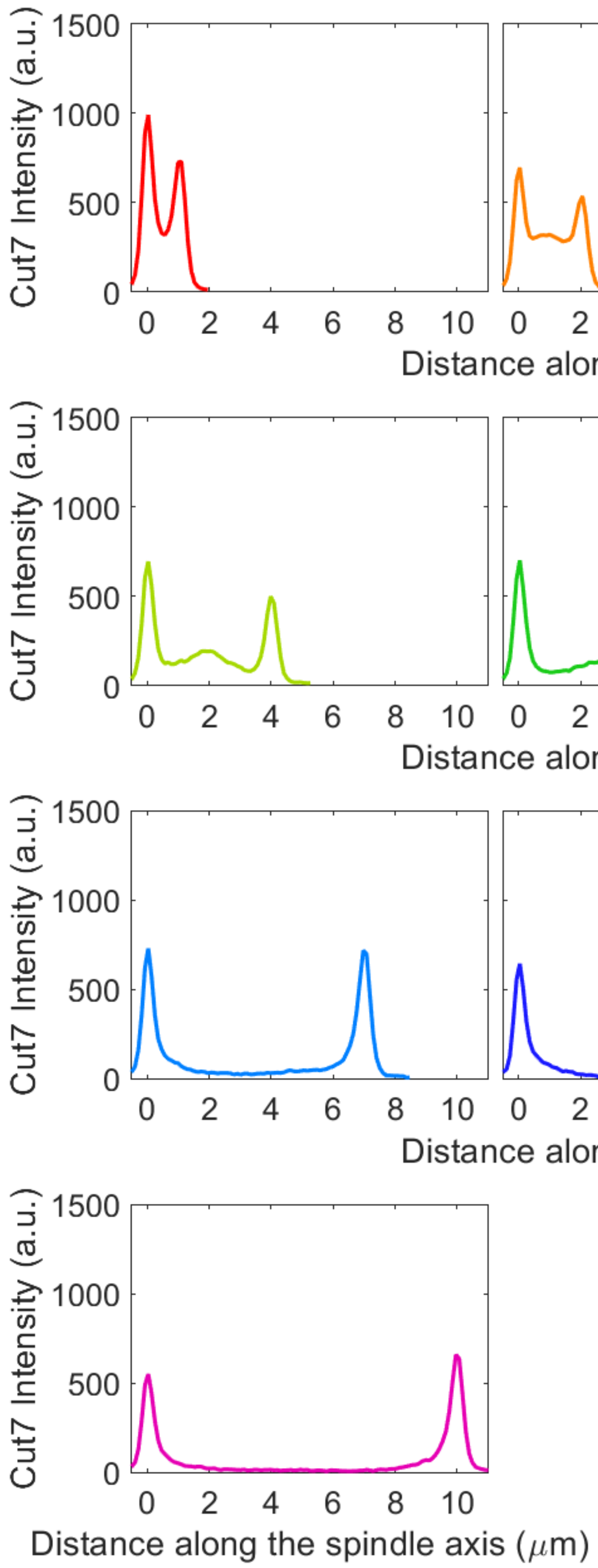


**Fig. S5.** Average Cut7-989-1028Δ-GFP intensity along the spindle axis for varying spindle length. Intensity data from 30 *cut7-989-1028Δ* cells.

| Strain | 37°growth | 37°spindle | 25°spindle length | Midzone-to-pole ratio | Interpretation |
|---|---|---|---|---|---|
| FL | ✓ | 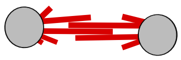 | ✓ | 1.0 | |
| 1-1032 | ✓ | 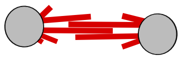 | ✓ | 1.5 | Sliding force↓ compensated by localization↑ |
| 1-1017 | ↓↓ | 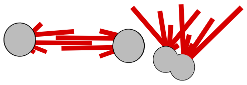 | ↓ | 1.4 | Sliding force↓↓ partially compensated by localization↑ |
| 989-1028Δ | ✗ | 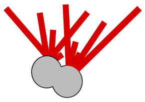 | ↓↓ | 0.9 | Sliding force↓↓ localization↓ severe defects |
| 1-1006 | ✗ | n/a | n/a | n/a | Lethal due to Cut7p loss of function |
| 1-988 | ✗ | n/a | n/a | n/a | |

**Fig. S6.** Summary of phenotypes of tail truncation mutants in the *pkl1+ klp2+* background, showing for each strain growth and spindle phenotypes at 37°C, pre-anaphase spindle length at 25°C, and average midzone-to-pole ratio.

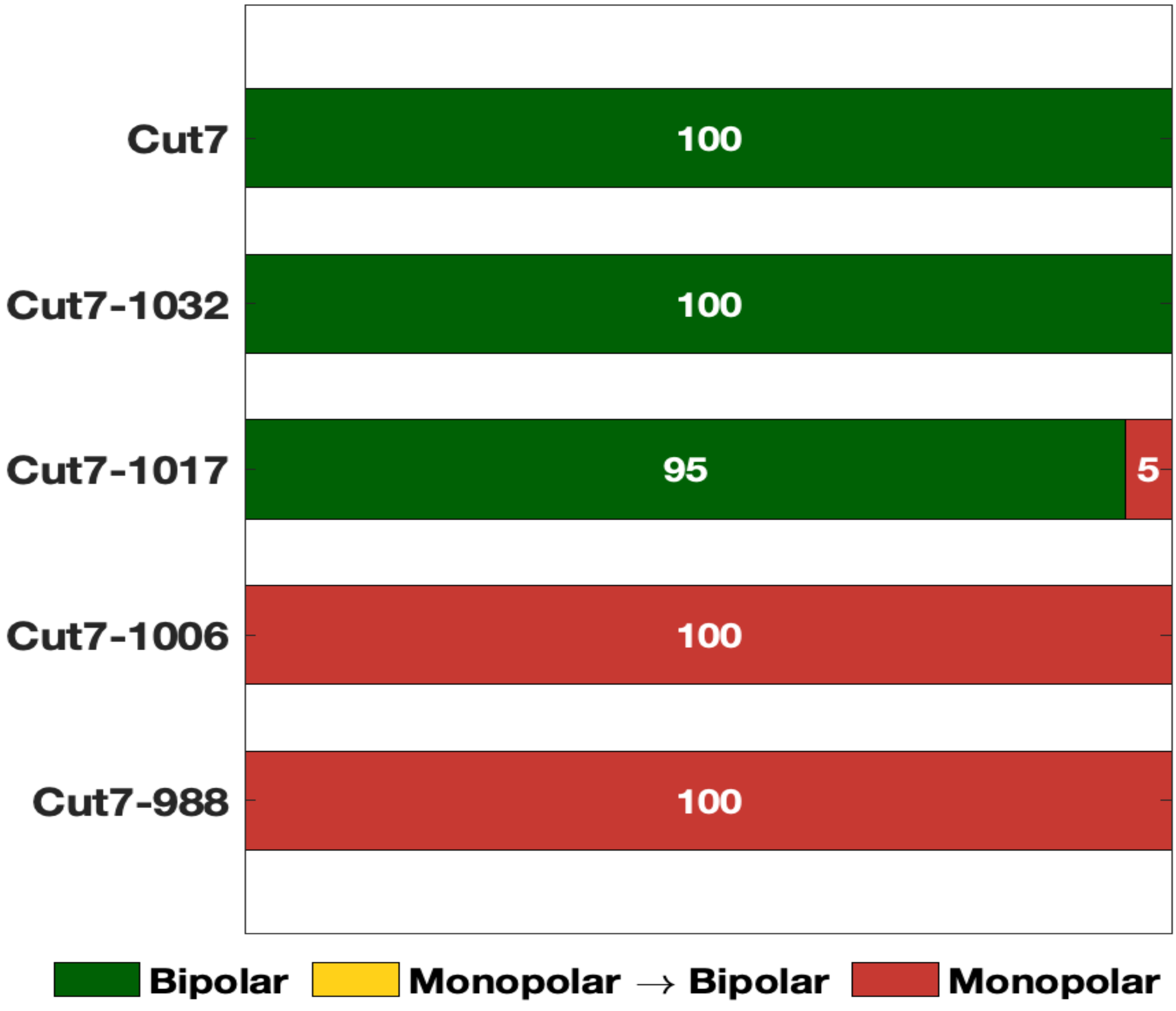


**Fig. S7.** Fraction of cells grown at 37°C that formed bipolar spindles (green), monopolar spindles that became bipolar (yellow), or persistent monopolar spindles (red) for 7 *cut7*, 21 *cut7-1032*, 21 *cut7-1017*, 15 *cut7-1006* and 16 *cut7-988* cells in the *pkl1*Δ*, klp2*Δ background.

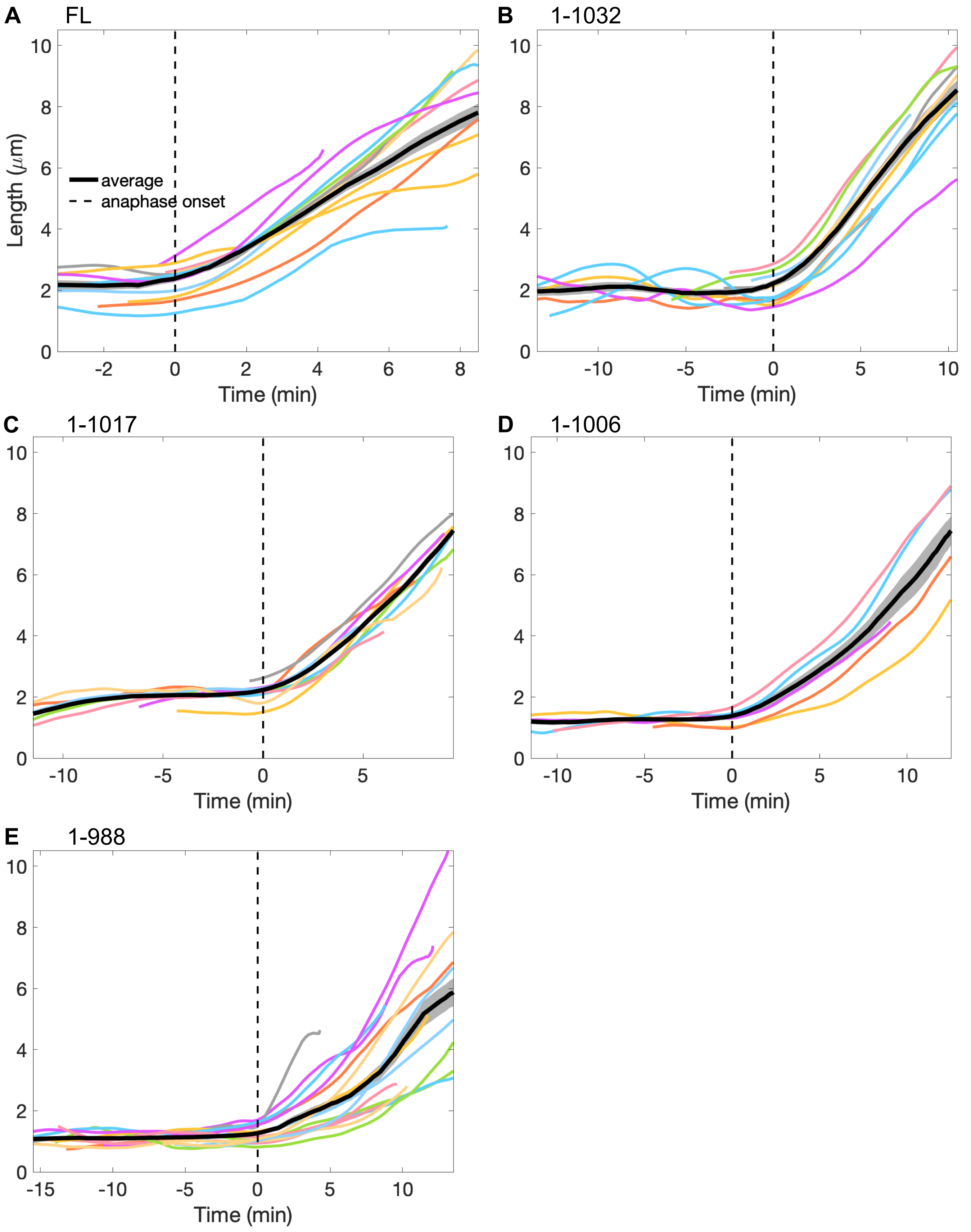


**Fig. S8.** Spindle length as a function of time showing curves from individual cells (colored lines) and average (black line). We defined $t = 0$ as the time of anaphase onset to align the curves (dashed line, Methods). Data acquired from 14 *cut7* (A), 13 *cut7-1032* (B), 11 *cut7-1017* (C), 6 *cut7-1006* (D), and 17 *cut7-988* cells (E) in the *pkl1*Δ*, klp2*Δ background. Only cells without visible protrusions were analyzed.

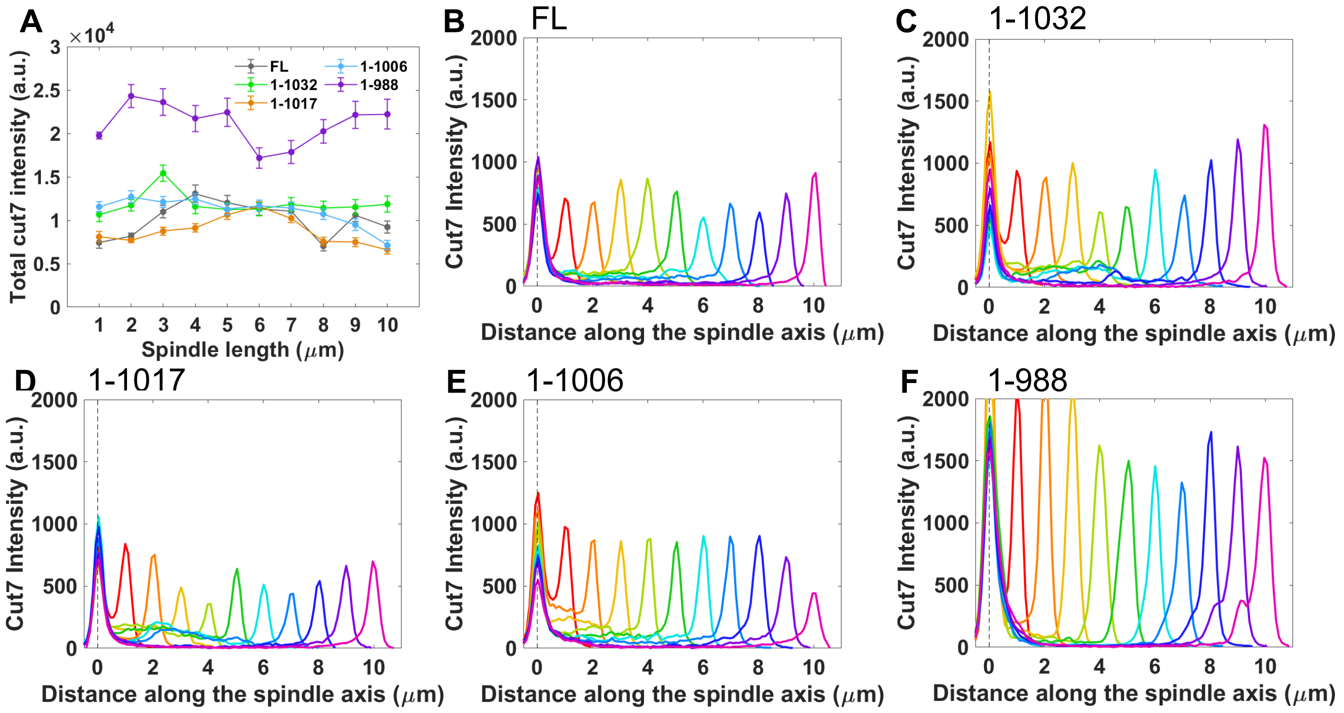


**Fig. S9.** (A) Total Cut7-GFP intensity on the spindle as a function of spindle length in the *pkl1*$\Delta$*, klp2*$\Delta$ background. Data from 44 *cut7*, 34 *cut7-1032*, 35 *cut7-1017*, 13 *cut7-1006*, and 40 *cut7-988* cells. (B-F) Average Cut7-GFP intensity along the spindle axis for Cut7-FL-GFP (B, from 44 cells), Cut7-1032-GFP (C, from 34 cells), Cut7-1017-GFP (D, from 35 cells), Cut7-1006-GFP (E, from 13 cells), and Cut7-988-GFP (F, from 40 cells).

| Strain (yMB) | Genotype | Figures |
|---|---|---|
| 1084 | pku80::hphMX6 z:adh15:mCherry-atb2:natMX6 ura4-D18 leu1-32 | - |
| 1149 | cut7-1-1085(FL)-GFP-kanr z:adh15:mCherry-atb2:natMX6 pku80::hphMX6 ura4-D18 leu1? his2? | Figure 2,3 |
| 1125 | cut7-1-1032-GFP:kanR pku80::hphMX6 z:adh15:mCherry-atb2:natMX6 ura4-D18 leu1-32 | Figure 2,3 |
| 1142 | cut7-1-1017-GFP:kanR pku80::hphMX6 z:adh15:mCherry-atb2:natMX6 ura4-D18 leu1-32 | Figure 2,3 |
| 1199 | cut7-full length-GFP (silent sites) pku80::hphMX6 z:adh15:mCherry-atb2:natMX6 ura4-D18 leu1-32 his2+ | - |
| 1168 | cut7(Δ989-1028)-GFP pku80::hphMX6 z:adh15:mCherry-atb2::natMX6 ura4-D18 leu1-32 | Figure 2,3 |
| 1091 | pku80::hphMX6 z:adh15:mCherry-atb2::natMX6 ura4-D18 leu1-32 his1::ura4+ | - |
| 1147 | pkl1D::his3+, klp2D::ura4+ z:adh15:mCherry-atb2:natMX6 leu1-32 his3- ura4-D18 | - |
| 1162 | cut7-1-1085(FL)-GFP-kanR, pkl1D::his3+, klp2D::ura4+ z:adh15:mCherry-atb2:natMX6 leu1-32 his3- ura4-D18 | Figure 4,5 |
| 1210 | cut7-1-1032-GFP:kanR, pkl1D::his3+ klp2D::ura4+ z:adh15:mCherry-atb2:natMX6 leu1-32 his3- ura4-D18 | Figure 4,5 |
| 1207 | cut7-1-1017-GFP:kanR, pkl1D::his3+ klp2D::ura4+ z:adh15:mCherry-atb2:natMX6 leu1-32 his3- ura4-D18 | Figure 4,5 |
| 1238 | cut7-1-1006-GFP:kanR, pkl1D::his3+, klp2D::ura4+ z:adh15:mCherry-atb2:natMX6 leu1-32 his3- ura4-D18 | Figure 4,5 |
| 1155 | cut7-1-988-GFP:kanR, pkl1D::his3+, klp2D::ura4+ z:adh15:mCherry-atb2:natMX6 leu1-32 his3- ura4-D18 | Figure 4,5 |



**Table S1.** ***S. pombe*** **strains used in this work**
